# Epigenetic characterization and manipulation of rare circadian neuron subtypes

**DOI:** 10.64898/2026.01.21.700891

**Authors:** Pranav Ojha, Gillian Berglund, Albert D. Yu, Shlesha Richhariya, Michael Rosbash

**Affiliations:** Howard Hughes Medical Institute, Dept. of Biology, Brandeis University, Waltham, MA 02454

## Abstract

The molecular circadian clock ticks away similarly in several distinct locations throughout the *Drosophila* head and body, yet regulation of this clock and specifically that of master transcription factor gene *Clock* (*Clk*) and its transcription is largely unknown. ATAC-Seq assays indicate that *Clk* chromatin appears inaccessible in most head neurons outside the circadian brain network, indicating that circadian clock functionality is gated by access to *Clk* chromatin. Moreover, distinct *Clk* enhancers function in circadian neurons versus glia, corresponding to distinct transcripts. To address the chromatin landscape of *Clk* with more temporal and spatial resolution, we developed a novel, high-purity, nuclear purification method, Elution-based INTACT (El-INTACT). It enabled the first time-resolved, six-timepoint chromatin accessibility landscape of ∼120 *Drosophila* circadian neurons/brain. It moreover allowed chromatin accessibility characterization of the 16 PDF-expressing ventrolateral circadian neurons (LNvs), which identified an additional, previously unidentified *Clk* enhancer. A CRISPR/Cas9 multiplexed gRNA-based strategy was used to disrupt these enhancers and indicated striking functional specificity. The findings establish a chromatin-level logic for cell type-specific circadian regulation and show that El-INTACT and the multiplexed gRNA-disruption of enhancer function are broadly applicable tools for exploring chromatin regulation in rare cell populations.

## Introduction

The circadian clock ticks away in many if not most eukaryotic organisms. It regulates a large fraction of gene expression, which cause the cycling of a myriad of physiological and behavioral functions (1–3). These endogenous gene expression patterns can anticipate the 24 fluctuations of light/dark cycles, temperature, nutrition, predators, potential mates among other environmental influences (4). The impact on such a diverse collection of genes and functions is because the circadian clock is keeping time in many if not most cells and tissues, which includes the mammalian brain (5). Nonetheless, in the mammalian brain, clock gene expression is probably strongest in the suprachiasmatic nucleus (SCN), the approximately 20,000 neurons in the anterior hypothalamus. Moreover, the SCN is known to play a crucial role in circadian behavior and physiology (6, 7). The fly brain has a similar region of very strong clock gene expression, the ca. 240 *Drosophila* circadian neurons (8–10). This small fraction of the fly brain is uniquely responsible for the generation of circadian behavioral rhythms (11, 12). There is evidence that both these regions, the SCN and the fly brain circadian neurons, are special for other reasons: for example, their anatomy and/or gene expression profiles are designed to interact with and respond to the external light-dark cycle (13–15) .

The molecular oscillator inside circadian neurons consists of a transcription-translation feedback loop (TTFL), which generates ∼24 hours periodicity (16–19). It has a similar architecture in mammals and flies, and most of its components are orthologs, i.e., it was inherited in both systems from a common ancestor (20, 21). In flies, the transcription factors CLOCK (CLK) and CYCLE (CYC) heterodimerize to activate transcription of target genes (19, 22, 23). These include *period* (*per*) and *timeless* (*tim*). Their protein products, PER and TIM, accumulate, translocate to the nucleus, repress CLK:CYC activity and eventually degrade, which closes the loop roughly every 24 hours (24, 25).

There is extensive information on the transcriptional regulation of *per* and *tim*. CLK: CYC directly activates these two genes by binding to their regulatory E-box enhancer elements (3, 26–28). Although this regulation is similar in different cell types, cell-type-preferential enhancers contributes to *tim* transcription: clock neurons but not glia use an intronic enhancer in addition to the common promoter enhancer (3).

We considered that a similar logic might apply to the gene *Clk*. This possibility is particularly intriguing because CLK:CYC occupies a central position in the circadian transcriptional network: the absence of *Clk* expression abolishes the broad transcriptional oscillations observed across the genome (29), and *Clk* expression in non-clock cells can generate ectopic clocks (3, 23, 30). Importantly, *Clk* has no established transcriptional regulatory elements analogous to E boxes for *per* and *tim* expression, leaving the cis-regulatory basis of its rhythmic expression essentially unknown.

To ameliorate this issue, we isolated different clock expressing and non-clock expressing nuclei from *Drosophila* heads and performed ATAC-Seq experiments to characterize the *Clk* chromatin landscape. The first experiments indicated that that *Clk* chromatin is inaccessible in most neurons outside the clock network, suggesting that the circadian clock is only present in a very limited fraction of the brain. A major feature of this paper is a new nuclear purification method Elution based-INTACT (El-INTACT). We improved the existing INTACT (31, 32) protocol by adding elution and FACS steps making it ideal for high purity, high recovery, and rapid processing of nuclei from genetically defined rare neuron populations. Using El-INACT, we isolated sufficiently pure circadian neuron chromatin to generate the first time-resolved chromatin accessibility landscape of the fly circadian network. El-INTACT also indicated that there is a novel enhancer enriched in the 16 PDF expressing circadian neurons. A second contribution is a CRISPR/Cas9/multiplexed gRNA enhancer disruption strategy. It shows that the *Clk* cis-regulatory architecture is highly cell type-specific. All of these results provided the first functional map of *Clk* chromatin and establish El-INTACT and CRISPR/Cas9-based enhancer disruption as broadly applicable tools for the profiling and manipulation of chromatin in rare cell populations.

## Results

### Accessibility of *Clk* chromatin is highly tissue-specific

We performed ATAC-Seq on broad populations of clock expressing and non-clock expressing sorted neurons from *Drosophila* heads and found that *Clk* chromatin displays striking cell type-specificity. The *Clk* cis-regulatory region contains three prominent accessible peaks, which we named according to the cell types in which they were identified: a neuron-specific peak, a common peak, and a glia-specific peak (Fig. 1A). The neuron-specific peak is accessible only in circadian neurons. The common peak is accessible in every other cell type known to harbor a molecular clock — circadian neurons, glia, eyes, and heads; the latter presumably mostly reflects a mixture of glia and eye chromatin. The glia-specific peak is so named because we first identified it in glia; it is present in eyes and heads as well as glia but is entirely absent from circadian neurons. Notably, a broad neuronal population (nSyb-Gal4) in which circadian neurons make up only a small fraction show no accessible peak at all within this region, despite a prominent peak in the adjacent gene (Fig. S1). This suggests that most and perhaps all head neurons outside the clock network lack the chromatin architecture required to express this master regulator, further suggesting a rather limited presence of molecular clocks within the brain (See Discussion.) A similar heterogeneous pattern of cell-type-specific enhancer usage is evident at other core circadian loci: *vri*, *pdp1*, and *cry* are each associated with distinct enhancer elements in different cell types (Fig. S1). Some of these enhancers probably contribute to the non-circadian expression of these genes outside of the clock network where they serve non-clock functions.

**Fig. 1.**
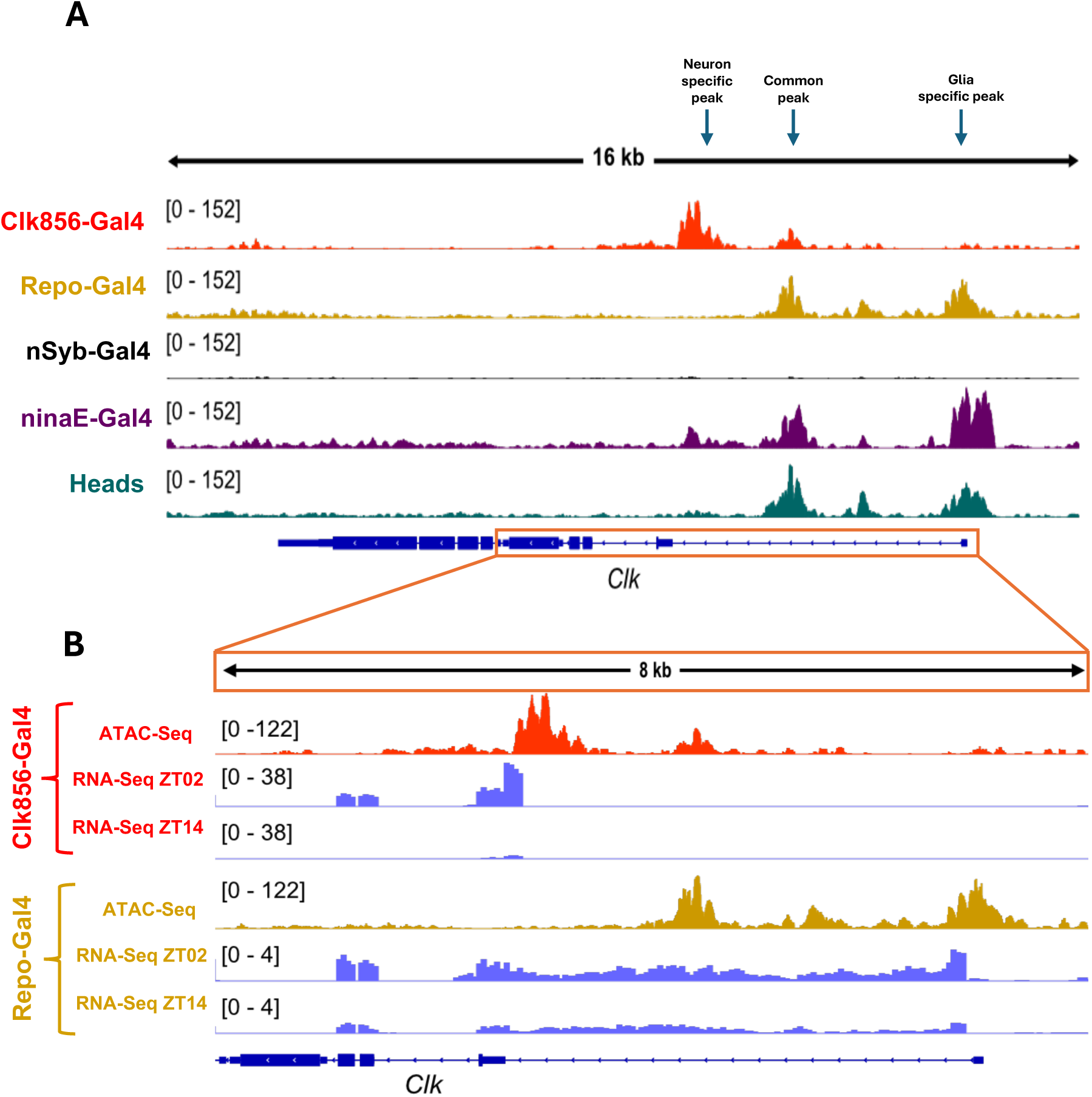
Accessibility of Clk chromatin is highly tissue-specific. A) Chromatin accessibility profiles for the *Clk* gene in clock neurons (Clk856-Gal4), glia (Repo-Gal4), all brain neurons (nSyb-Gal4), photoreceptor neurons (ninaE-Gal4), and heads. We identified 3 distinct peaks in these cell types: neuron-specific, common and glia specific peak. All ATAC-Seq profiles were generated from flies collected at ZT02. B) Zoomed-in version of the *Clk* cis-acting region with ATAC-Seq and corresponding RNA-Seq profiles collected at two timepoints – ZT02 and ZT14.

Might the neuron and glial peaks have direct functional consequences for *Clk* transcription? Closer inspection and comparisons with RNA-Seq profiles (Fig. 1B) indicate that the transcription start sites differ dramatically between circadian neurons and glia, and that both RNA profiles cycle robustly.

#### El-INTACT reliably purifies low copy number nuclei from fly heads

To capture the dynamics of the accessible peaks in circadian neurons and within rare circadian neuron subpopulations, we developed a new method called Elution-based INTACT, which greatly improved the clean isolation of neurons of interest. El-INTACT is a 3-step nuclear purification strategy - Enrich, Elute, and FACS-purify (Fig. 2).

**Fig. 2.**
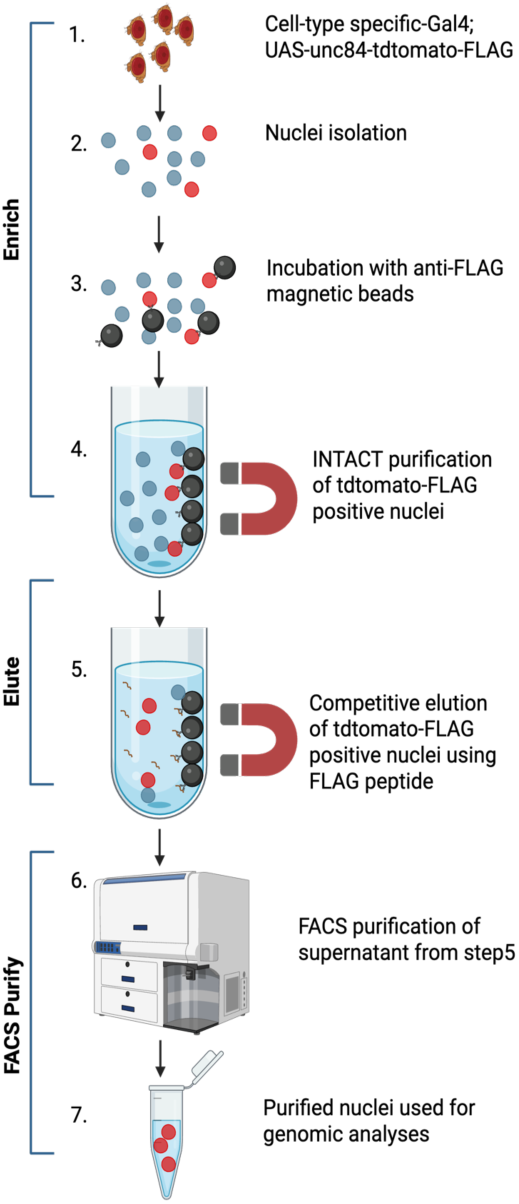
Schematic of El-INTACT. Elution based INTACT (El-INTACT) is a 3-step method – enrich, elute, and FACS purify. Enrich: Steps 1-4. Heads from flies expressing UAS-unc84-tdtomato-FLAG driven by a cell-type specific-Gal4 are isolated and homogenized using a mini-INTACT homogenizer (31). This homogenate is a mixture of all head nuclei, including specific nuclei expressing unc84-tdomato-FLAG on their nuclear membrane. Nuclei of interest are enriched by incubating the homogenate with anti-FLAG magnetic beads. 2) Elute: Step 5. By adding FLAG peptide, nuclei bound to anti-FLAG are competitively eluted off the beads. Although these nuclei are enriched in the eluted homogenate, many sticky non-tdtomato-FLAG nuclei remain bound. 3) FACS purify: Steps 6-7. Nuclei of interest are finally purified by sorting based on tdtomato fluorescence.

To provide an initial characterization of El-INTACT, we utilized our workhorse Clk856-Gal4 driver (Fig. S2A) that expresses in ∼120 fly brain circadian neurons (9, 33). When nuclei are sorted directly after brain or head homogenization, circadian neurons make up ∼ 0.1 – 0.01% of the total nuclei population; this estimate is based on about 140,000 brain neurons and 1 million fly head cells. In contrast, El-INTACT purified circadian neurons are ∼10% of the total nuclear population, indicating a dramatic enrichment of between 100X and 1000X (Fig. S2B). As a result, the sorting rate is also substantially enhanced.

#### ATAC-Seq peaks around the clock

Several studies have examined time-series transcriptomics datasets from *Drosophila* circadian neurons (2, 34). However, there is no comparable time-series chromatin accessibility profile from the same source. To this end, we used El-INTACT to isolate circadian nuclei from 6-time points (ZT02, ZT06, ZT10, ZT14, ZT18, ZT24) and performed ATAC-Seq. Many ATAC-Seq peaks undergo robust circadian cycling, and these five canonical circadian genes (*per*, *tim*, *vri*, *pdp1*, *Clk*) are among the top genes based on cycling amplitude (Fig. 3A-F). Further, most of the top cyclers are in two classes: early morning (ZT02) cyclers and early night (ZT14 and ZT18) cyclers (Fig. 3E).

**Fig. 3.**
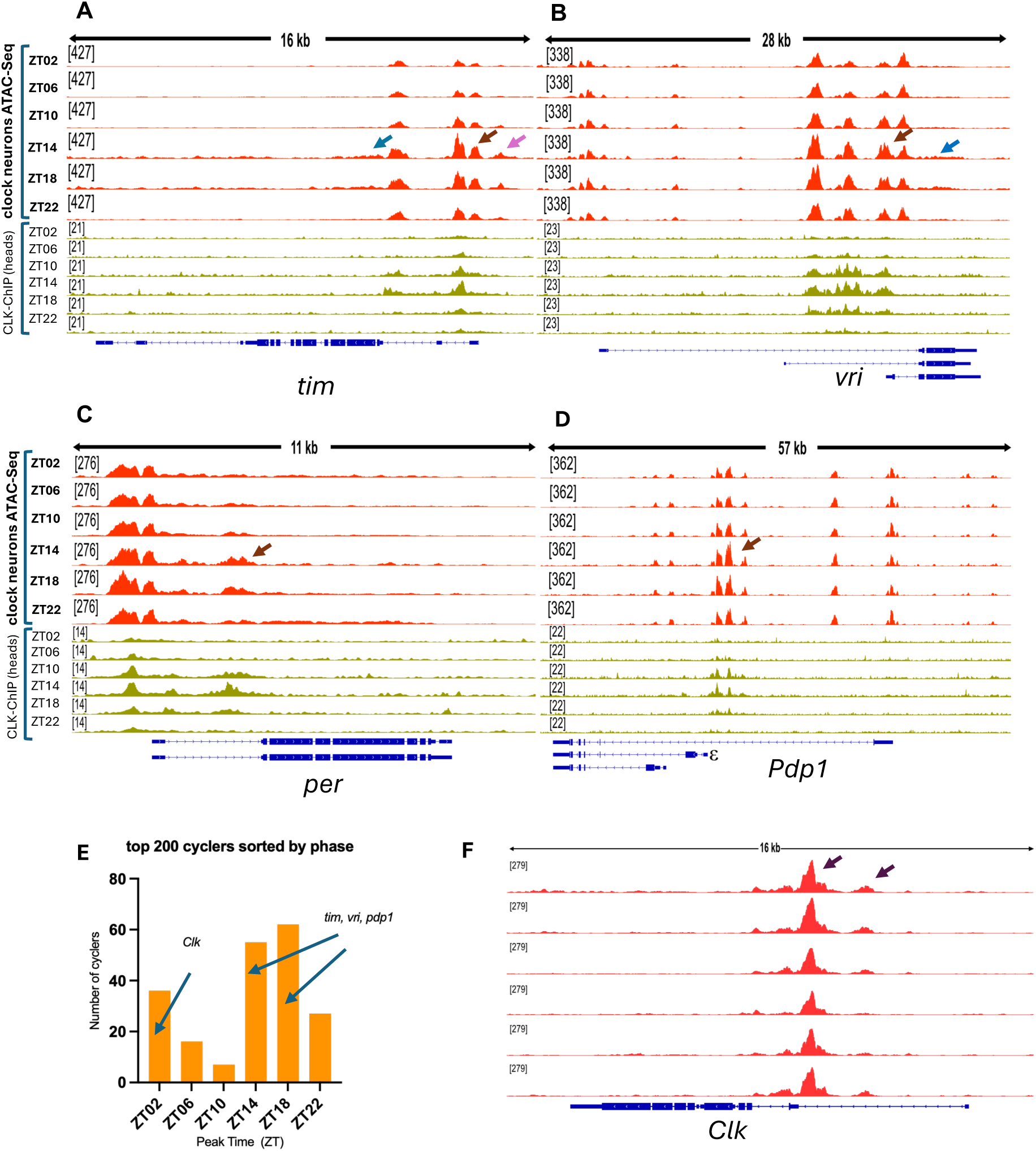
El-INTACT driven 6 time point ATAC-Seq from the clock neurons reveals distinct cycling patterns in core circadian genes. In red are 6 time-point ATAC-Seq profiles of core circadian genes and in mustard green are 6 time-point CLK-ChIP profiles from heads for the same circadian genes. CLK-ChIP dataset was directly taken from a previous publication (3). Here, brown arrows indicate cycling of the chromatin directly overlapping with ChIP-Seq, pink arrow indicates the cycling of adjacent chromatin and blue arrows indicate chromatin in the gene body. (A and B) *tim* and *vri* chromatin cycle with a peak at ZT14. C) The intronic peak in *per* cycles robustly in line with CLK-ChIP cycling. D) Only the circadian isoform of PDP1 (*Pdp1*ε) cycles within clock neurons. E) top 200 cycling ATAC-seq peaks (based on their JTK values) cluster into two major groups, ZT02 cyclers and ZT14/18 cyclers. F) Both peaks in the *Clk* chromatin cycle with a peak cycling at ZT02. Both ATAC-Seq and CLK-ChIP profiles have at least two replicates.

Visually inspecting the four core circadian genes that are direct targets of CLK:CYC indicate that their cycling ATAC-seq peaks are coincident with their CLK:CYC binding peaks (Fig. 3, indicated by brown arrows). For example and consistent with previous reports (3, 35), the promoter as well as the intronic *tim* ATAC-Seq peaks cycle robustly (Fig. 3A). Interestingly, there are also adjacent accessible regions within *tim* that are not direct CLK:CYC targets but also cycle robustly (Fig. 3A, indicated by pink arrow). This same pattern occurs with another top circadian gene cycler *vri*: there are CLK:CYC as well as adjacent cycling peaks (Fig. 3B). We suggest that this adjacent cycling is influenced by heightened CLK:CYC binding during peak transcription, ZT14-18 for *tim* and *vri*. In addition, chromatin accessibility within the gene body (blue arrows) is cycling. This presumably reflects Pol II and transcriptional cycling (see Discussion).

#### El-INTACT ATAC-Seq identifies enhancers from even rarer neurons

Some rare circadian neuron subtypes like the l-LNvs (large ventrolateral neurons), are difficult to isolate via the traditional brain dissection method followed by FACS sorting; this is probably because of their large size and complex projection patterns (Fig. S4). In any case, genomic RNA-seq libraries from sorted circadian neurons, both bulk and single-cell datasets, have a dramatic underrepresentation of l-LNvs (36). Because El-INTACT purifies nuclei, it should not have this same issue. We therefore compared ATAC-Seq profiles of circadian nuclei purified using El-INTACT with those purified using standard brain dissection + FACS-sorting. Chromatin for genes specific or highly enriched in l-LNvs like *Pdf* (9, 37) and *dimm* (38), were both much better represented in El-INTACT purified clock nuclei (Fig. S3), indicating that El-INTACT is indeed superior in characterizing chromatin from these difficult-to-isolate cells.

#### Purification of LNv nuclei and identification of an additional *Clock* enhancer

Directly FACS sorting LNv nuclei from fly heads without any enrichment has unacceptable issues including a slow sorting rate. (See Discussion). This results in an insufficient number of nuclei for clean and robust ATAC-Seq profiles which is ∼4000 (Fig. 4A). We therefore expressed the UAS-Unc84-tdtomato-FLAG nuclear tag under PDF-Gal4 control, which only expresses in the 16 PDF-expressing-LNv cells. El-INTACT not only gives a 10-fold higher yield of nuclei but also dramatically reduces the hands-on time of dissection + FACS sorting (Fig. 4A).

**Fig. 4.**
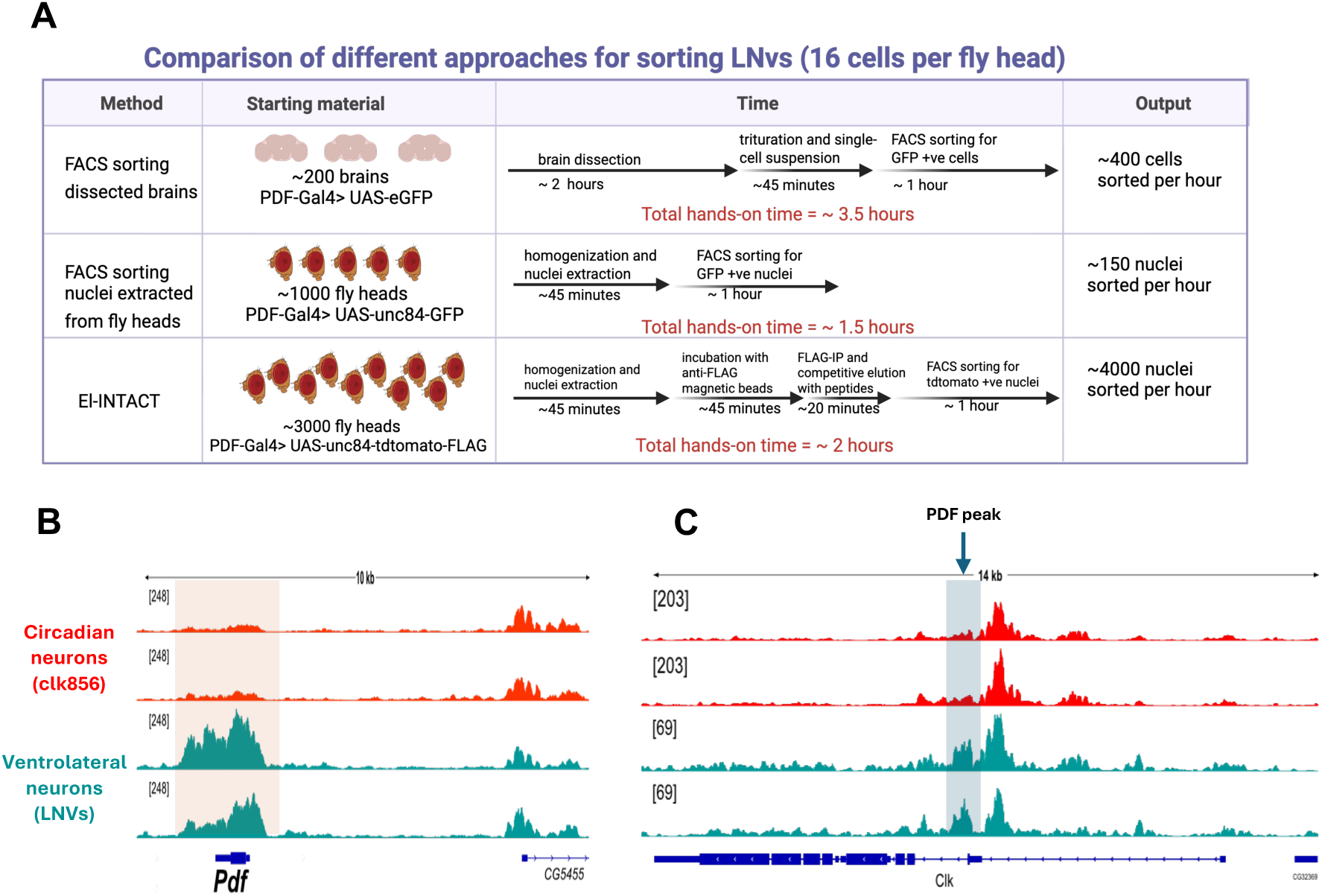
Isolation of LNvs using El-INTACT and PDF specific *Clk* cis acting peak. A) To compare different purification approaches for the 16 LNvs per fly head, first approximately 200 PDF-Gal4>UAS-eGFP brains were dissected, triturated into single cell suspension, and sorted for 1 hour. This process required around 3.5 hours of hands-on time and yielded ∼400 LNv neurons. Second, 1000 PDF-Gal4> UAS-unc84-GFP fly heads were homogenized, their nuclei extracted and sorted for 1 hour. Fly heads make for a significantly dirtier homogenate than dissected brains, and therefore FACS sorting LNv nuclei from this prep lead to a much lower yield compared to dissection and FACS sorting. El-INTACT required 3000 PDF-Gal4>UAS-unc84-tdtomato-FLAG fly heads as the starting material and yielded 4000 nuclei in an hour of sorting. Flies used for El-INTACT-ATAC-Seq were collected and frozen at ZT02. B) ATAC-Seq profile of the gene *Pdf* is more accessible in El-INTACT isolated LNv nuclei (teal) than El-INTACT isolated Clk856 nuclei (red). This accessibility profile is in line with the gene-expression pattern of *Pdf* in the LNvs (9). C) ATAC-Seq profile of the gene *Clk* in PDF neurons has a unique peak called PDF peak, indicated by blue arrow. 4A was created in Biorender. Ojha, P. (2025).

To confirm the successful purification of LNv nuclei, we compared the *Pdf* gene ATAC-Seq profile from LNv nuclei with the same profile from all circadian (Clk856) nuclei. The LNv *Pdf* profile shows significantly greater chromatin accessibility within the gene body, indicating successful LNv nuclear purification (Fig. 4B). Moreover, this LNv-specific *Clk* profile appears to contain an additional regulatory element. We named it the PDF peak, and it is immediately adjacent to the neuron-specific peak described above (Fig. 4C). We note that the Y-axis scale differs between the two profiles, most likely because the Clk856-Gal4 enhancer element itself overlaps the neuron-specific peak, inflating its apparent size relative to the profiles generated using a different Gal4 driver. Despite this scale difference, the size of the PDF peak relative to the neuron-specific peak in the LNv profile strongly suggests that this newly identified region reflects a genuine, distinct regulatory element.

#### CRISPR-Cas9 mutagenesis identifies a functional enhancer

The above results indicate that the *Clk* cis-regulatory region contains at least 4, largely cell-type-specific accessible elements: a PDF peak, a neuron-specific peak, a common peak, and a glia-specific peak. Although we do not know which proteins occupy those peaks, one or more of these elements might contribute to *Clk* transcription and circadian behavior. To address this possibility, we generated a panel of fly lines using the CRISPR/Cas9 system to mutagenize individual peaks with multiplexed guide RNAs (22). Each line carries gRNAs against an individual accessible region (Fig. S4A), and we also generated a positive-control line targeting the *Clk* coding sequence directly (Table S2). As in our previous studies using CRISPR/Cas9 to disrupt protein function in circadian neurons (23, 34), the multiplexed gRNAs and Cas9 were both under UAS control, allowing us to restrict disruption to specific cell populations using the appropriate Gal4 driver. As a further positive control, we assayed a deletion of the entire *Clk* cis-regulatory region (ClkOUT); as expected, this rendered flies arrhythmic and additionally disrupted the dorsal projections of clock neurons (Fig. 5A, S4B) (39).

**Fig. 5.**
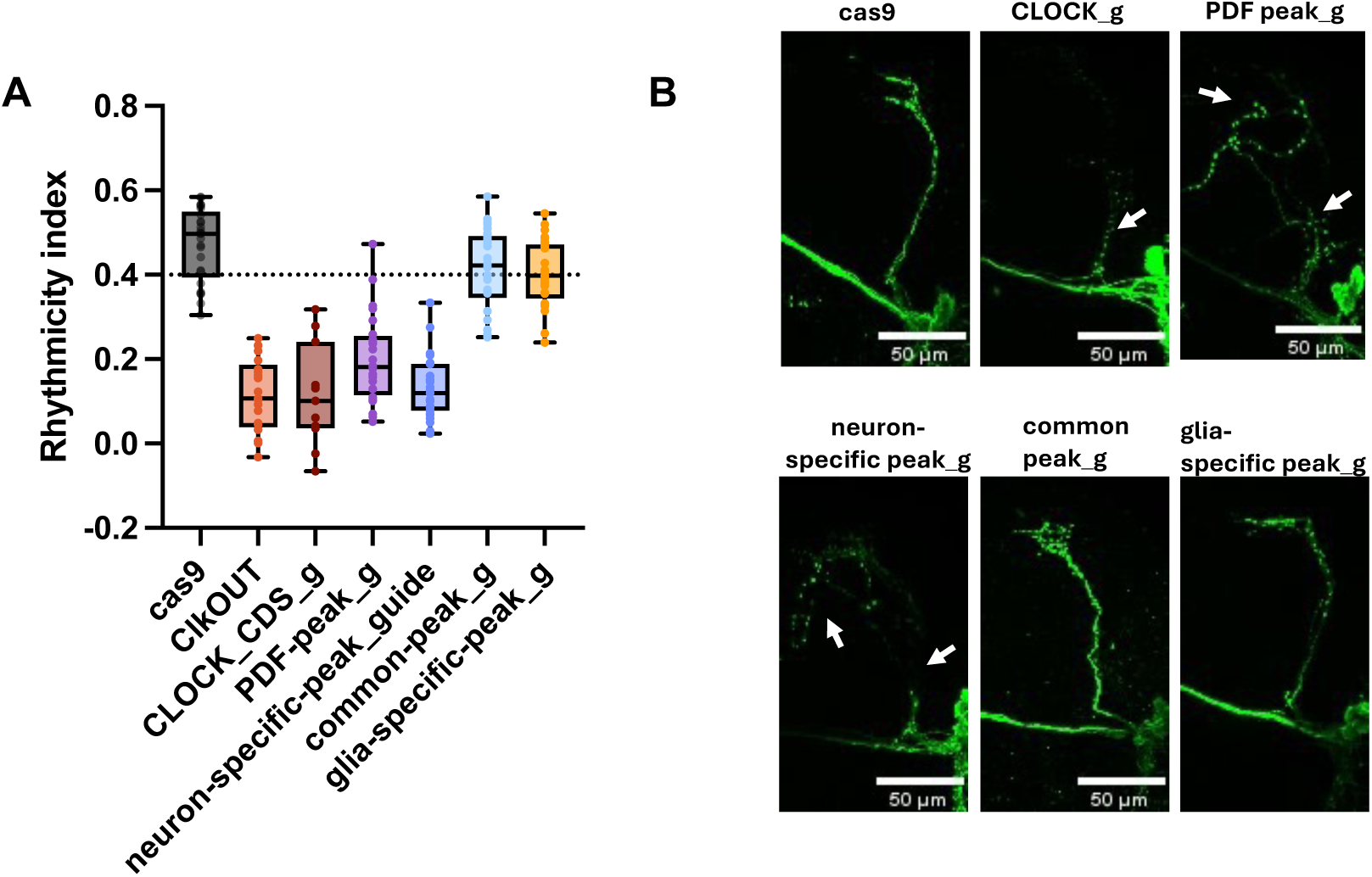
CRISPR based disruption of El-INTACT identified *Clk* enhancers. A) Rhythmicity index, an indicator of circadian rhythm strength of ∼10-day-old flies with indicated mutations driven by elav-Gal4. B) Imaging of the sLNV projections in elav-Gal4 x guide mutants using anti-PDF staining. Disrupted and missing projections are indicated by white arrows.

To generate deletions comparable to the germ line, whole body deletion in ClkOUT, we used elav-Gal4 to drive Cas9 and peak-specific gRNAs pan-neuronally. We confirmed successful disruption of these targeted regions by PCR as previously shown (Fig. S8) (40).

Disrupting the common peak or the glial peak had no detectable effect on circadian behavior (Fig. 5A), perhaps because the common peak disruption did not decrease *Clk* expression sufficiently and the glial clock does not contribute to circadian behavior (41). In contrast, disrupting either the PDF peak or the neuron-specific peak alone was sufficient to render flies arrhythmic by standard locomotor activity criteria, but disrupting the neuron-specific peak alone produced a stronger phenotype than disrupting the PDF peak alone (Fig. 5A). Consistent with this observation was the morphology of the PDF neurons dorsal projections: disrupting the common peak or the glia-specific peak line was indistinguishable from the control, whereas disrupting the *Clk* coding sequence directly, the neuron-specific peak, or the PDF peak affected the projections to varying degrees (Fig. 5B, S4). Moreover, combining disruption of the neuron-specific peak with either the common or the glia-specific peak produced phenotypes indistinguishable from disruption of the neuron-specific peak alone, consistent with the interpretation that the common and glia-specific peak contribute little to *Clk* expression (Fig. S9). Together, these results indicate that the neuron-specific and PDF peaks but not the common or glia-specific peaks have a functional role in orchestrating *Clk* transcription and rhythmic circadian behavior.

#### Molecular consequence of peak disruptions

If disruption of the neuron-specific and PDF peaks compromises *Clk* transcription, this should be reflected in altered expression of direct CLK targets. We first examined peak PER protein levels by immunostaining at ZT02, focusing initially on the four large LNvs (l-LNvs) per hemisphere (Fig. 6). Disruption of the PDF peak eliminated PER staining in approximately 25–50% of l-LNvs (Fig. S6), whereas disruption of the neuron-specific peak eliminated staining in 50–75% of l-LNvs compared to the normal 4 l-LNvs in elav-Gal4>Cas9 control flies (Fig. 6A). This corresponds to roughly two to three affected cells per hemisphere for the neuron-specific disruption. Unaffected l-LNvs were indistinguishable from control l-LNvs (Fig. 6A). Disruption of the common and glia-specific peaks had no effect on PER staining (Fig. S9).

**Fig. 6.**
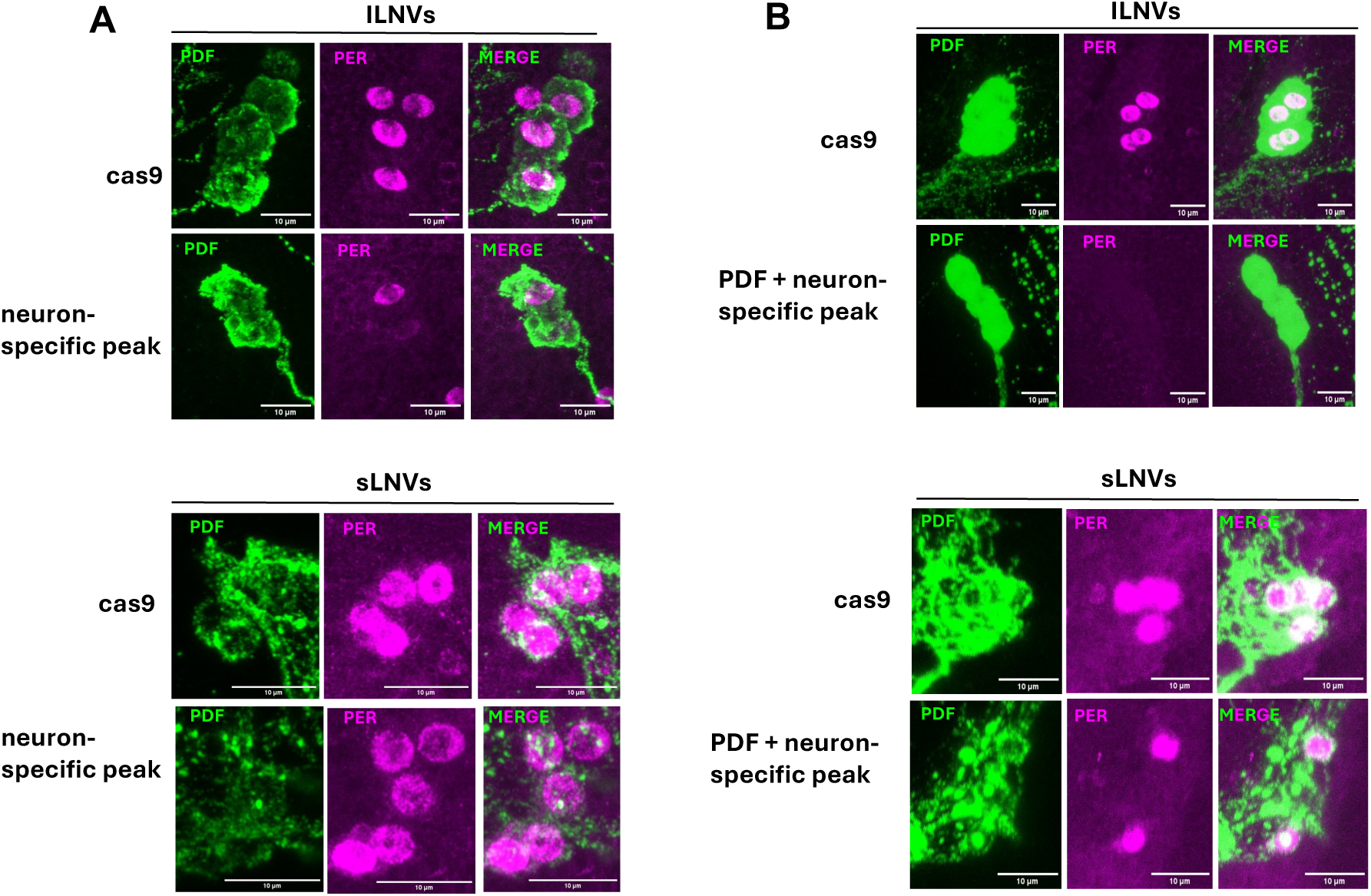
PER staining large and small LNVs in different CRISPR mutants. A) PER staining (magenta) in large LNVs, unlike in small LNVs, is vulnerable to neuron-specific peak disruption. Large and small LNVs are identified by staining with anti-PDF (green). B) Combing the neuron specific peak disruption with PDF peak disruption results in a complete loss of PER from large LNVs and a 50% loss in PER from small LNVs.

We also examined the small LNvs (s-LNvs) under the same conditions. Unexpectedly, they were essentially unaffected by any single-peak disruption. Because other clock neuron clusters such as the DN1ps showed the same sensitivity pattern as the l-LNvs (Fig. S5), the s-LNvs appear to be the outliers. Notably, combining gRNAs against the PDF and neuron-specific peaks together eliminated PER staining in essentially 100% of l-LNvs and 50% of s-LNvs (Fig. 6B). The data suggest that PER expression is more robust in s-LNvs than in l-LNvs or in other clock neurons.

To assess transcription more directly, we performed *tim* in situ hybridization at ZT14, when *tim* expression normally peaks (3). Single disruption of the PDF peak, the neuron-specific peak, or the glia-specific peak were very similar to the PER expression result (Fig. 7A). Combined disruption of the PDF-specific and neuron-specific peaks abolished *tim* expression in essentially all l-LNvs (Fig. 7B) and s-LNvs (Fig. 7C). Although these data are somewhat different than the PER staining results directly above and suggest CLK:CYC-driven transcription of *per* and *tim* is not completely identical, most features of the disruptions are similar. Perhaps some of the neuron type differences simply reflect more *Clk* expression in s-LNvs than l-LNvs.

**Fig. 7.**
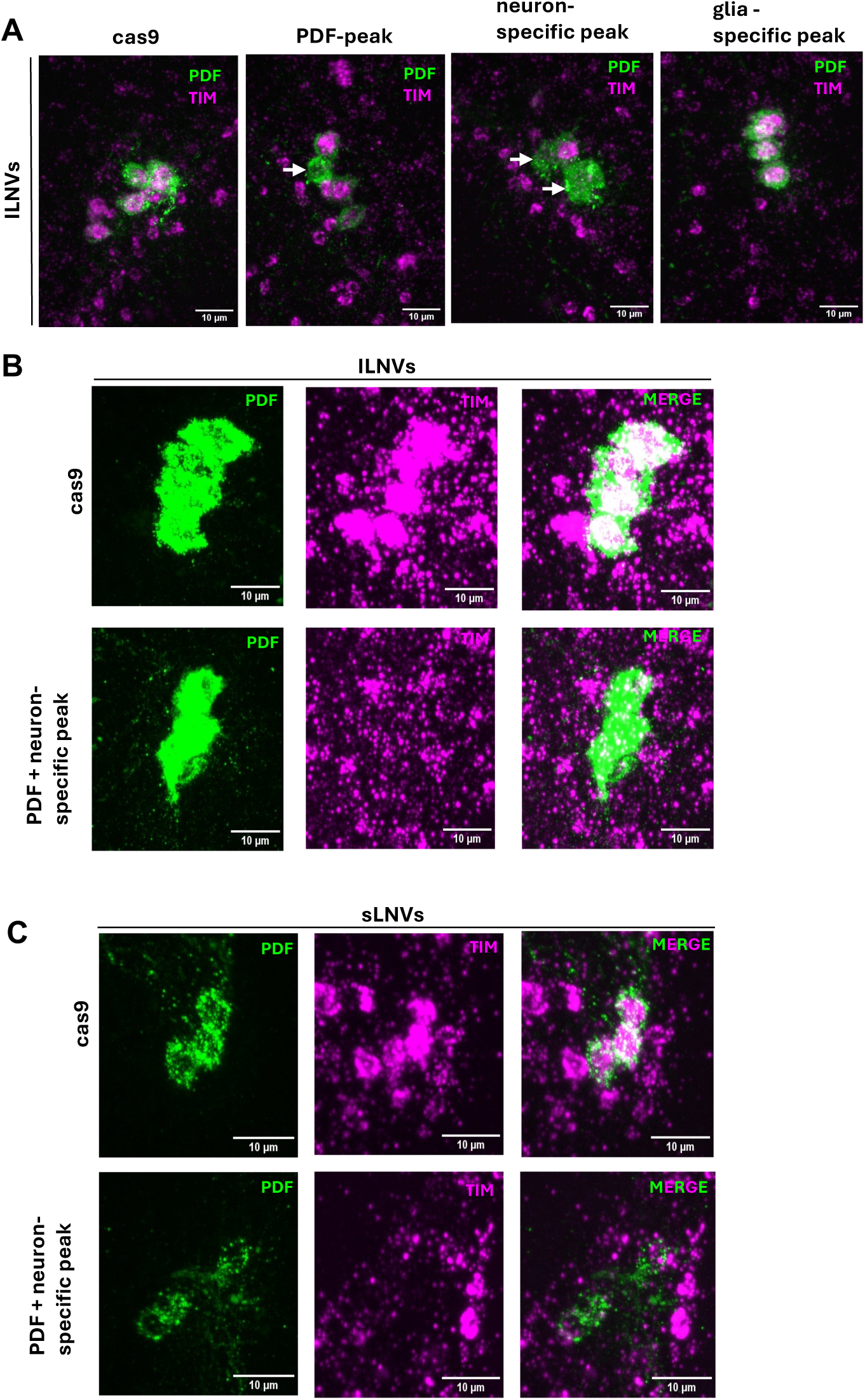
*timeless* in-situ in large and small LNVs in different CRISPR mutants. A) *timeless* in-situ (magenta) in cas9 control and individual peak disrupted flies. Large LNVs are identified by staining with anti-PDF (green). White arrows indicate large cells missing *tim* expression. B & C) Combing the neuron specific peak disruption with PDF peak disruption results in a complete loss of *tim* from large and small LNVs.

## Discussion

The work in this paper describes two technical advances to purify and manipulate in vivo chromatin of rare, genetically defined cells: El-INTACT and CRISPR-based enhancer disruptions. We applied these methods to the adult fly brain and especially to its ca. 120 clock neurons and one of its subpopulations, and we focused on the regulatory chromatin that impacts the expression of the master circadian regulator gene *Clk*. Despite the simplicity and common features of the circadian system wherever it is expressed, the regulation of *Clk* is surprisingly heterogeneous within the brain and especially within this circumscribed set of neurons.

### Insights from and advantages of El-INTACT

Many, and perhaps most, adult *Drosophila* central brain neurons exist in only a handful of copies per brain (42, 43). Combined with the rich collection of Gal4 driver lines marking discrete neuronal subsets, El-INTACT can efficiently purify nuclei from precise low-copy-number populations. We recently exploited this approach for RNA-seq profiling and show here it also generates exceptionally clean ATAC-Seq profiles from ∼120 circadian neurons per brain. This enabled an informative time course of cycling ATAC-Seq peaks, some of which are located within known genes under circadian transcriptional control. We then used El-INTACT to purify and characterize the16 PDF-expressing clock neurons per brain, a population essentially inaccessible to standard approaches.

There are several advantages of El-INTACT compared to conventional dissection and trituration: First, purified nuclei can be used directly for downstream molecular assays without additional centrifugation steps that lose material; second, frozen heads can be banked and pooled as needed, adding logistical flexibility; third, the speed of the frozen head protocol and its large amount of starting material avoids both the labor-intensive dissection and the lengthy FACS sorting times that limit the more standard approaches. This frozen head protocol and large amount of starting material provide a fourth advantage: it is essential to generate enough nuclei to achieve high resolution profiles of rare cell types.

Our experience is that 3000-4000 *Drosophila* nuclei reliably yields high-quality, replicable ATAC-Seq profiles. This bulk-profiling strategy also overcomes the issues that plague single-nuclei chromatin profiling; its signal: noise is good enough for cell-type identification but often insufficient for more detailed insights (44, 45). In contrast, El-INTACT-enabled bulk ATAC-Seq profiling allows the signal aggregation from thousands of pooled nuclei and thereby circumvents this missing-data problem. This consideration may be particularly important for rare *Drosophila* neurons, for which the compact fly genome likely enhances the sparsity constraints that even mammalian chromatin profiling face with single-nuclei methods.

Moreover, brain dissection and trituration is known to be problematic for another reason, namely, specific cell types can be missing (46). This problem is well-illustrated by our own prior 10X circadian neuron data: several known clock neurons present in the fly brain at 2-4 cells/hemisphere were nearly absent from the 10X datasets compared to other clock neurons similarly present identically in vivo (9, 10, 47). These almost missing neuron types included the large PDF-expressing LNvs (l-LNvs) and two evening cell types (trissin- and ITP-expressing LNds), almost certainly because the large cell bodies and extensive processes of these neurosecretory neurons are poorly recovered by dissection and FACS. This is likely why there has been no profiling of these otherwise well-studied l-LNvs (9, 36) in the 15 years since they were first hand-sorted for bulk RNA-Seq (38, 48). El-INTACT recovered these neurons in their expected numbers.

The reliable and clean ATAC-Seq profiles in our 6 time point dataset identified many features including cycling ATAC-seq peaks (Fig. 3). Those that correspond to CLK-CYC binding to core clock gene regulatory elements within *per* and *tim* genes had been previously described, but this was from a 2 time point dataset with a quite modest differences between peak and trough values (35). Our dataset identifies these same peaks but with much bigger cycling amplitudes, as much as 2-to 4-fold, which likely reflect the much higher signal:noise ratio afforded by El-INTACT. There are in addition cycling enhancer peaks adjacent to the CLK-CYC peaks. These unknown peaks have the same temporal profiles as CLK-CYC binding, suggesting that this known pioneer transcription factor (49) may loosen histone-DNA contacts and help recruit additional factors to associate with adjacent chromatin.

There was also cycling open chromatin within the gene bodies of the core clock genes (Fig. 3, blue arrows). They peaks during times of maximum transcription, suggesting that they most likely reflects actively transcribing polymerases.

The cycling ATAC-Seq dataset also identified many additional peaks with impressive amplitudes; like the CLK:CYC peaks, they most likely reflect the temporal association of other transcription factors or chromatin proteins with circadian regulatory elements, which probably contribute to enhancing or repressing nearby transcription in a circadian manner. A challenge for the future is to identify the protein factors responsible for these and other orphan ATAC-Seq peaks.

Finally, the ATAC-Seq dataset identified putative cell type-specific circadian enhancers. We focused on those within *Clk*, in part because its regulation had not been well-explored. These striking cell type-specific enhancers suggest a strategy in which individual cell types establish their own cis-regulatory solutions for gene expression rather than relying on a universal enhancer architecture. This strategy may reflect the highly cell type-specific nature of transcription factor expression in discrete neuron populations and is also in agreement with our previous cell type-specific characterization of *tim* chromatin (3). Moreover, this strategy appears more general as it also applies to other core clock genes like *vri*, *pdp1* and *cry*; they have their own cell type-specific enhancer patterns (Fig. S1).

### Cis-regulatory control of *Clk* transcription

The distinct cis-regulatory strategy of *Clk* is most striking in the simple circadian neuron versus glia profiles; they result in two distinct transcripts arising from distinct transcription start sites (Fig. 1B). These transcripts are the well-annotated isoform in glia, and a shorter, previously unidentified isoform specific to circadian neurons, likely previously unidentified because it is masked in bulk tissue by the dominant glial signal. This difference in 5′UTR length could plausibly affect transcript stability and/or translation independent of transcription rate, and RNA differences track with each population. Glial *Clk* transcription rises to a broad peak around ZT20 that persists until ZT02, whereas neuronal *Clk* transcription peaks sharply at ZT02 and declines rapidly thereafter, cycling dynamics that are based on our own and previously published single-cell data (9, 47, 50). Although these features warrant further investigation, the shorter neuronal transcript might support faster turnover and thus a sharper, more tightly tuned transcriptional cycle; it might even be important for coherent behavioral rhythms.

The complete absence of accessible *Clk* chromatin in non-clock neurons (Fig. 1, nSyb-Gal4) indicates that a simple but profound regulatory layer restricts which neurons can run a canonical molecular clock. Because overexpressing CLK in adult neurons is sufficient to open chromatin at several of its target genes (3), this regulation is very different from a downstream failure to respond to CLK activity. The closed chromatin at *Clk* may therefore restrict the circadian transcriptional program to the 240 circadian neurons within the 100,000 fly central brain neurons. There is little comparable information for the mammalian brain.

### Insights from CRISPR-based enhancer disruptions

To functionally assess candidate enhancers within *Clk*, we used cell type-specific Cas9 expression along with cell type-specific multiplexed guide RNA expression (23, 24, 34). This strategy builds on our previous work of generating targeted germ line deletions to disrupt individual E-box sites within *tim* regulatory DNA (3). The same approach was not possible for *Clk* as it lacks characterized regulatory motifs. We therefore used multiplexed guide RNAs to disrupt individual candidate regulatory regions and assess their functional relevance based on ATAC-Seq data independent of motif content. This approach acknowledges that an ATAC-Seq peak may reflect protein binding to chromatin without an effect on gene expression.

We chose this multiplexed CRISPR/Cas9 approach over CRISPRi. The latter is effective in cell culture and mammalian systems but was ineffective against the same *Clk* cis-regulatory targets in flies; dCas9-KRAB produced no phenotype. Moreover, CRISPRi has not been shown to have detailed enhancer-specific resolution, suggesting multiplexed CRISPR/Cas9 mutagenesis may be a more broadly useful tool.

As in our previous protein-coding knockout work (40), multiplexed guides produced heterogeneous, PCR-validated disruptions (Fig. S8). Although this might produce mosaic outcomes, the reproducible differences as well as the additive effect of combining guides support a genuine difference in enhancer function between l-LNvs and s-LNvs rather than a repair difference between cell types. We suggest that some of the enhancer function differences parallel other distinctions between the two LNv subtypes. Only the s-LNvs are the master pacemaker cells and retain robust transcriptional oscillations even in constant darkness. Moreover, s-LNvs are born far earlier in development (L1 larvae) than l-LNvs (pupae). Strikingly, CLK is required for s-LNv survival: s-LNvs are largely missing or malformed in ClkOUT flies, whereas l-LNvs develop normally in the same genetic background. This suggests that the enhancer disruption results might reflect effects on s-LNv specification as well as s-LNv target gene expression but only effects on target gene expression in l-LNvs.

### Extending El-INTACT to other organisms

Both methods should generalize well beyond this study. El-INTACT should be applicable to any *Drosophila* tissue or cell type, including entirely uncharacterized rare populations in the central complex or antenna. It should also extend to other insects, especially mosquitoes, butterflies and silkmoths as cell-type-specific genetic tools mature in these species (39–42). The enhancers identified this way may in turn aid construction of more specific driver lines in these non-model organisms. El-INTACT should also translate to mammalian tissue and cell culture, requiring only prior transgenic delivery of unc84-tdTomato-FLAG; this is achievable via viral methods with high cell-type specificity). Moreover, its compatibility with frozen starting material makes it well suited to diverse genomic and epigenomic profiling of mammalian cells and tissues.

## Methods

### Drosophila Lines

Flies were housed in standard cornmeal/agar medium with yeast under 12:12 hrs LD (light:dark) cycles. UAS-unc-84-tdtomato-FLAG (31) flies were a generous gift from Sean Eddy. Other fly strains used in this study are in Appendix, Table S1. CRISPR lines expressing tandem gRNA under UAS-control were generated using previously described protocols (51). CRISPR Optimal Target Finder tool (52) was used to identify gRNA specific to target peaks. These guides were custom synthesized and incorporated into a pCFD6 vector (Addgene #73915) by GenScript (Piscataway, New Jersey). Plasmids were injected into embryos by Rainbow Transgenic Flies Inc (Camarillo, CA, USA). Positive transformants were screened for by red eye color based on the mini-white marker.

### Circadian Activity Assay

Circadian behavior was assayed as described previously (53). Briefly, one week old male flies were used for the behavioral experiments, and locomotor activity of individual flies was measured using *Drosophila* activity monitors (Trikintetics Inc), in which individual flies were placed into glass tubes with food (2% agar and 4% sucrose) on one end and a plug to close the tube on the other end. DD behavior from flies were recorded and analyzed from day 2 to day 7 of DD using Sleep and Circadian Analysis MATLAB Program (SCAMP) (54). Each experiment was performed twice and got similar results.

### Immunohistochemistry

Immunohistochemistry for *Drosophila* brains were performed as described previously (55). Briefly, flies were fixed in 4%PFA (Fisher Scientific #50-980-487) then brains were dissected and blocked in blocking buffer (10% Normal Goat Serum (NGS, Jackson Labs #005-000-121) in 0.5% PBST). The following primary antibodies in blocking buffer were used: anti-RFP (1:1000; Proteintech #5f8), rabbit anti-PER (Rosbash lab, 1:1,000), mouse anti-PDF (DSHB-PDF C7; 1:1000). Primary antibody incubations were done for 15-18h at 4°C. Secondary antibodies were used at 1:500 dilution in blocking buffer and were incubated at 4°C overnight. Brains were mounted in Vectashield (Vector Laboratories #H-1000-10) and imaged on Leica Stellaris 8 confocal microscope. Image J was used for signal quantification (56).

### RNA Scope

Fluorescence in situ hybridization was performed on *Drosophila* brains using the RNAScope Multiplex Fluorescent Detection Kit v2 (ACD) as previously described (3). Flies were collected at ZT14 and fixed whole in 4% formaldehyde/PBST (PBS + 0.5% Triton X-100) for 5h at 4 °C, then brains were dissected and fixed overnight under the same conditions. Brains were washed in PBST and PBST+1% BSA, then subjected to target retrieval (5.5 min at 100 °C in 1× Target Retrieval Solution), followed by sequential washes in PBST+BSA and methanol. Brains were post-fixed in 4% formaldehyde/PBST, washed, and treated with Protease Plus (10 min, 40 °C) before overnight hybridization with the Tim-C1 probe at 40 °C.

Signal amplification followed the manufacturer’s protocol, with all 40 °C incubations performed in a heat block and washes done 2×5 min in RNAScope Wash Buffer. The Tim-C1 probe was labeled with Opal 650 dye (1:6,000 in TSA buffer; 30 min at 40 °C).

For combined immunohistochemistry, brains were washed post-HRP blocking, blocked in 10% NGS (2 h RT or overnight 4 °C, protected from light), and incubated with mouse anti-Repo primary antibody (1:100, 24–48 h, 4 °C), followed by washes and Alexa Fluor 488 goat-anti-mouse secondary (1:200, overnight, 4 °C). Brains were washed and mounted in Vectashield. Imaging was performed on a Leica Stellaris 8 confocal microscope (40× oil immersion).

### FACS Sorting of cells and nuclei

Clock neurons labelled by Clk856-Gal4 and UAS-eGFP were purified using Fluorescence-activated cell sorting (FACS) using methods described previously (3). Clk856-Gal4>UAS-eGFP brains were dissected in Schneider’s media (SM, Gibco #21720001) and incubated with 0.75 mg/mL Collagenase (Sigma-Aldrich, #C1639) and 0.4 mg/mL Dispase (Sigma-Aldrich, #D4818) at room temperature for 30 min. Brains were then triturated 50 to 80 times to obtain a single-cell suspension for FACS. Hoechst stain (Invitrogen, #R37605) was used to identify live cells. GFP-positive cells of interest were collected with gates that were set using a sample without GFP. The same protocol was applied for sorting PDF-Gal4>UAS-eGFP flies.

FACS sorting of PDF-Gal4 expressing nuclei was enabled by crossing PDF-Gal4 flies to UAS-unc84-GFP flies (31). Fly heads isolated with vigorous vortexing followed by separation over dry ice cooled sieves. 1000 PDF-Gal4> UAS-unc84-GFP fly heads were added to 5 mL of homogenization buffer (250 mM sucrose, 1M Tris (pH 8.0), 1M KCl, 1M MgCL 2,10% Triton-X 100, 0.1M DTT, 1X complete protease inhibitor cocktail). After 20 tractions of the mini-INTACT homogenizer (57) at 1000 rpm, nuclei were filtered through 10 µm CellTrics strainer (Sysmex: 04-004-2324) and spun down at 800 g for 10 minutes at 4°C. Nuclei were resuspended in resuspension buffer (1% BSA in PBS with 0.1% Tween-20). Resuspended nuclei were filtered through a 35μm cell strainer into a 5 mL round bottom polystyrene FACS tube. Hoechst stain (Invitrogen, #R37605) was used to identify live nuclei. GFP-positive nuclei of interest were collected with gates that were set using a sample without GFP.

### El-INTACT

Fly heads were isolated with vigorous vortexing followed by separation over dry ice cooled sieves. 1000-3000 fly heads were added to 5 mL of homogenization buffer (250 mM sucrose, 1M Tris (pH 8.0), 1M KCl, 1M MgCL 2,10% Triton-X 100, 0.1M DTT, 1X complete protease inhibitor cocktail). After 20 tractions of the mini-INTACT homogenizer at 1000 rpm, nuclei were filtered through 10 µm CellTrics strainer (Sysmex: 04-004-2324) and spun down at 800 g for 10 minutes at 4°C. Nuclei were resuspended in resuspension buffer (1% BSA in PBS with 0.1% Tween-20) and incubated with 40 μl of anti-FLAG magnetic beads (FisherSci, PIA36797) for 45 minutes at 4°C with constant end-over-end agitation. Bead-bound nuclei were then collected on a magnetic rack and washed once with 500uL of resuspension buffer. Bead-bound nuclei were then resuspended in 80 μl of resuspension buffer.

For the elution step, 1mg of 3XFlag-peptide (Proteintech fp) was diluted in 100uL of resuspension buffer (1% BSA in PBS with 0.1% Tween-20). 20uL of the diluted peptide was added to 80 μl resuspended bead-bound nuclei and mixed gently. Flag-tagged nuclei were competitively eluted off the beads with gentle shaking on a thermomixer for 10 minutes at 600 rpm. Flag beads were collected on a magnet and the supernatant containing isolated nuclei was filtered through a 35μm cell strainer into a 5 mL round bottom polystyrene FACS tube. Hoechst dye (one drop per 0.5 ml of sample; Invitrogen, #R37605) was added into the sample tube to stain the nucleus. All steps were performed at 4°C. A BD Melody FACS machine in single cell sorting mode was used for cell collection. tdTomato- and Hoechst-positive single cells were collected in a 1.5-ml Eppendorf tube containing 0.5 ml of ATAC-Resuspension Buffer (RSB) (58) and used for downstream ATAC-Seq library preparation.

For RNA-Seq, all steps were performed at 4°C and RNase inhibitors were added to homogenization and resuspension buffers at 0.5% concentration. Importantly, the elution of nuclei with FLAG peptides requires using buffers without these inhibitors due to the presence of a reducing agent (DTT in RNAsin) that interferes with the peptide elution step and substantially reduces nuclei yield.

### Omni-ATAC-Seq

Transposition on sorted cells or nuclei was performed as previously described (3, 58). Briefly, cells sorted into 0.5 ml of ATAC-Resuspension Buffer were centrifuged at 500 rcf for 10 min at 4 °C. 50 µL of cold ATAC-Resuspension Buffer (RSB) containing 0.1% NP40, 0.1% Tween20, and 0.01% Digitonin was added to the pellet and incubated on ice for 3 minutes. Lysed cells were washed out by adding 1ml of cold ATAC-RSB containing 0.1% Tween-20 and inverted 3 times to mix. Nuclei were separated from lysed cells by centrifugation at 1000 rcf for 10min at 4 °C. Obtained nuclei pellet was used for ATAC-Seq.

FACS sorted nuclei in ATAC-RSB were centrifuged at 1000 rcf for 10min at 4 °C. Nuclei pellet was resuspended in 50 µL of Transposition mix (25 µL 2× TD buffer, 16.5 µL PBS, 0.05 µL 10% v/v Tween, 0.05 μL 1% v/v Digitonin,and 2.5 µL TDE1 Enzyme (Illumina, San Diego, CA, USA Catalog #20034198) and incubated at 37 °C for 30 min. Tagmented DNA was purified using Zymo DNA clean and concentrator kit. Purified DNA fragments were amplified using single-indexed Nextera primers (Integrated DNA Technologies) for 12 PCR cycles. Amplified libraries were purified using AMPure XP beads (Beckman Coulter, Brea, CA, USA Catalog #A63880), and size distribution was accessed using TapeStation High-Sensitivity D1000 Screentape. Final libraries were sequenced on a Nextseq 1000 (Illumina).

### ATAC-Seq Preprocessing

ATAC-Seq (fastq) files were adaptor-trimmed using fastp. Bowtie2 was used to align trimmed files using the following parameters: --local --very-sensitive-local --no-unal --no-mixed --no-discordant --phred33 -I 10 -X 700 (59). Samtools was used to remove PCR duplicates and Sambamba was used to remove multimapping reads (60). Tn5 insertion bias was corrected using a custom python script and peaks were called using MACS2.

### ATAC-Seq Differential Peak Calling and Normalization

Peaks called from MACS2 were intersected between replicates using bedtools, and common peaks found in at least 3 timepoints were merged and converted into a reference annotation in SAF format (61). Featurecounts was used to count reads in each sample within each region in the reference file and input into DESeq2 in R for between-samples normalization.

For circadian analysis, a normalized count table was used through the Meta2D function in MetaCycle and normalization factors were used to produce normalized bigwig files for visualization using Deeptools bamcoverage (62, 63). A Meta2D p-value threshold of 0.05 was used to define cycling peaks. Cycling peaks were putatively assigned to their target gene using Uropa (64). Identified cyclers were further analyzed using Nitecap (65).

### Smart-Seq3

PolyA-selected mRNA was isolated using the Dynabeads mRNA Direct Kit (Thermo Fisher #61011) and eluted in 9 µl water. Full-length cDNA with unique molecular identifiers (UMIs) was generated from 4.5 µl of this RNA using an adapted Smart-seq3 protocol. The original single-cell Smart-seq3 method (using Maxima H Minus Reverse Transcriptase, Thermo Fisher #EP0751) was scaled for bulk neurons by increasing all reaction volumes 5-fold and using 15 PCR cycles. Amplified cDNA was purified with 0.6× AMPure XP beads (Beckman Coulter) and assessed for quality and concentration via High Sensitivity D5000 ScreenTape (Agilent #5067-5592) on a TapeStation.

Approximately 750 pg of cDNA was used as input for library preparation with the Nextera XT DNA Library Preparation Kit (Illumina #FC-131-1096), modified to avoid overly small fragment sizes. Tagmentation was performed using 0.8 µl Amplicon Tagment Mix with 9.2 µl cDNA, 2× tagmentation buffer, and water, at 55 °C for 10 min, then quenched with 2.5 µl NT buffer. i5/i7 indexes (2.5 µl each) and 7.5 µl NPM were added on ice to a final volume of 25 µl, followed by 9 PCR cycles. Final libraries were purified with 0.7× AMPure XP beads (#A63881) and quantified via High Sensitivity D5000 ScreenTape on a TapeStation.

Libraries were sequenced on an Illumina NovaSeq 6000 at Novogene Corporation Inc. (Sacramento, CA, USA), yielding 20–38 million raw paired-end reads (2×150 bp) per sample.

## Data, Materials, and Software Availability

**Table S1.** List of fly strains used.

| Genotype | Description | Source |
| --- | --- | --- |
| Clk856-Gal4 | Gal4 labeling most circadian neurons | BDSC 93198 |
| nSyb-Gal4 | Gal4 labeling all neurons | BDSC 51635 |
| PDF-Gal4 | Gal4 labeling LNVs | BDSC 6899 |
| Elav-Gal4 | Gal4 labeling all neurons (on Chr X) | BDSC 458 |
| UAS-Cas9.P2 | Cas9 under UAS control (on Chr 2) | BDSC 58986 |
| UAS-neuron-specific-peak-guide | 4x tandem guides against the neuron specific peak | This study |
| UAS-common-peak-guide | 3x tandem guides against the common peak | This study |
| UAS-glia peak guide | 3x tandem guides against the glia specific peak | This study |
| UAS-CLOCK-CDS guide | 4x tandem guides against the clock CDS | This study |
| UAS-PDF peak guide | 3x tandem guides against the pdf peak | This study |
| 10xUAS-unc84-tdtomato-FLAG | Unc84-tdtomato-FLAG under UAS control (on Chr 2) | Gift from Sean Eddy (31) |
| UAS-eGFP | eGFP under UAS control | BDSC 5430 |

**Table S2.**
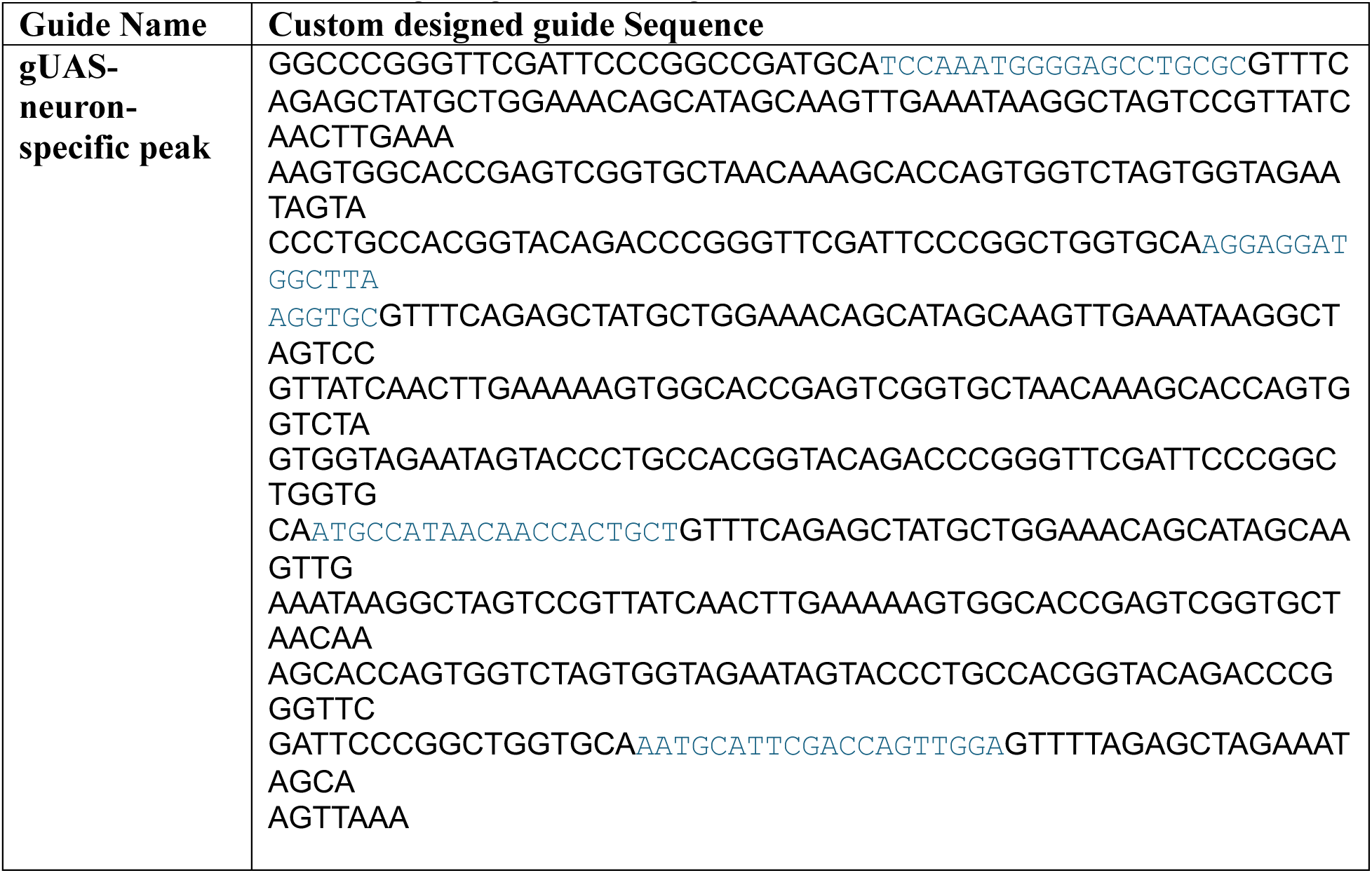
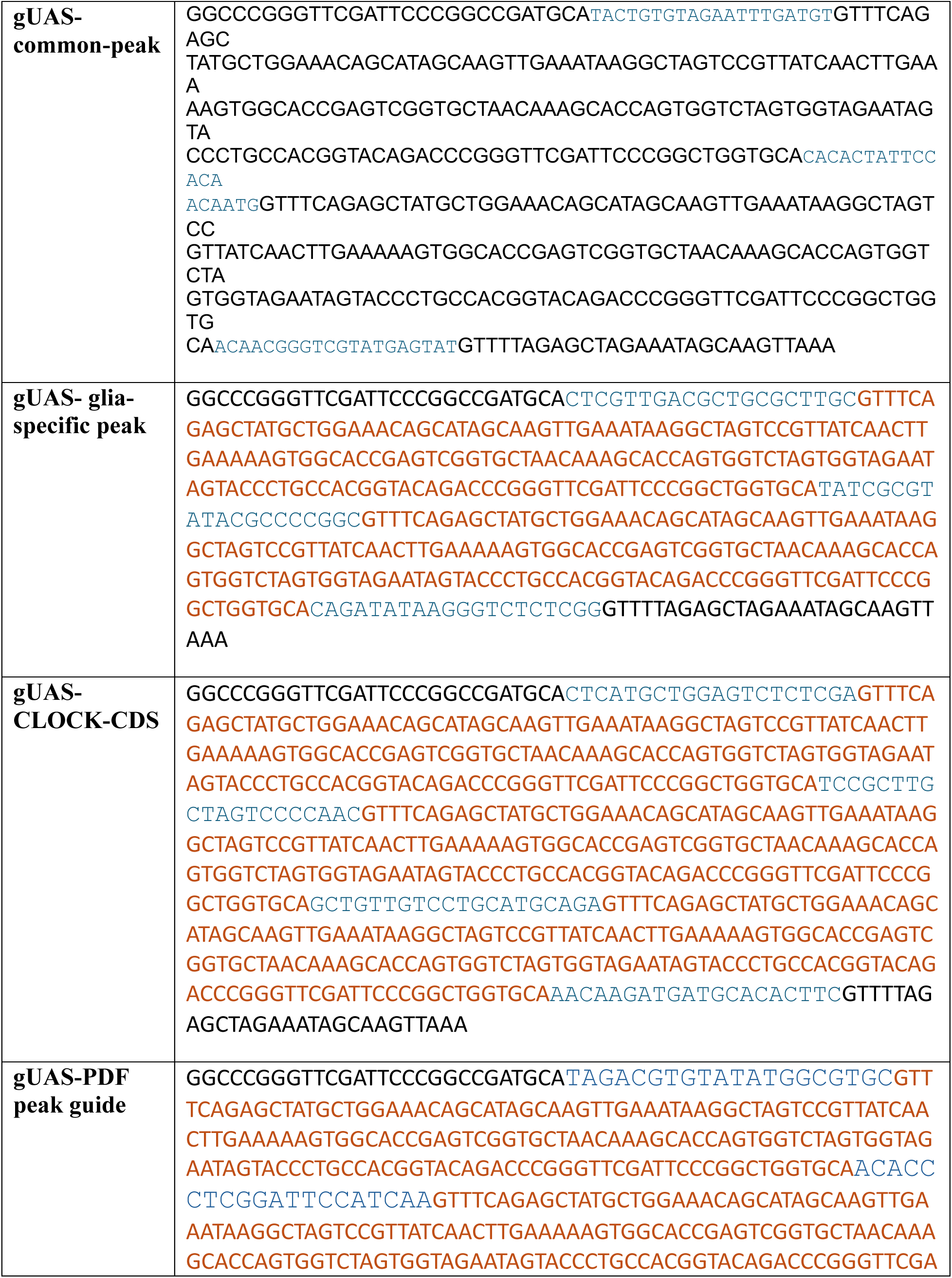
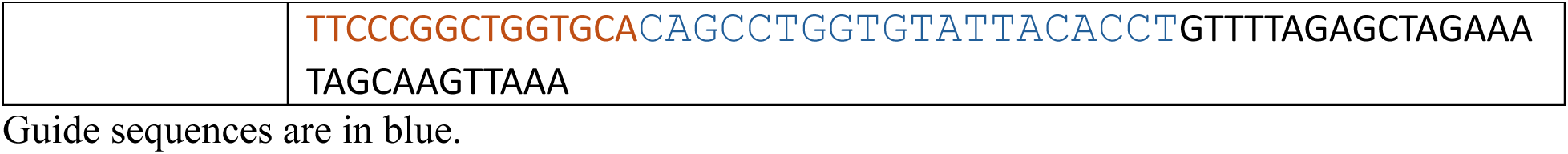
**List of custom designed guide RNAs against *Clk***

**S1.**
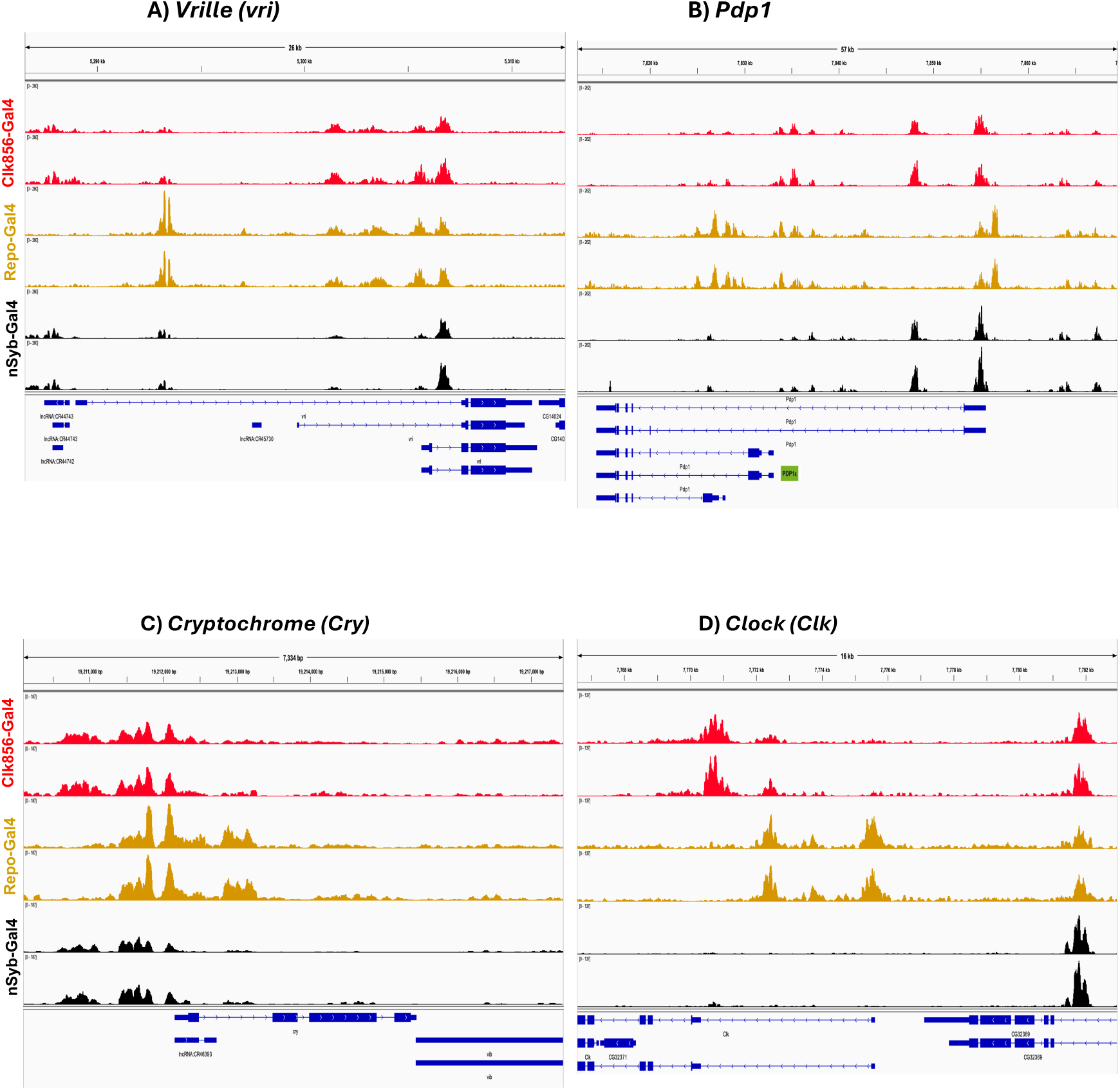
Cell-type specific differences in chromatin accessibility of circadian genes. Chromatin accessibility profiles of four circadian genes in sorted clock neurons (red), glial cells (orange), and all brain neurons (black). All cells were collected at ZT02. A) *Vri*, B) *Pdp1*, C) *Cry* and D) *Clock*

**S2.**
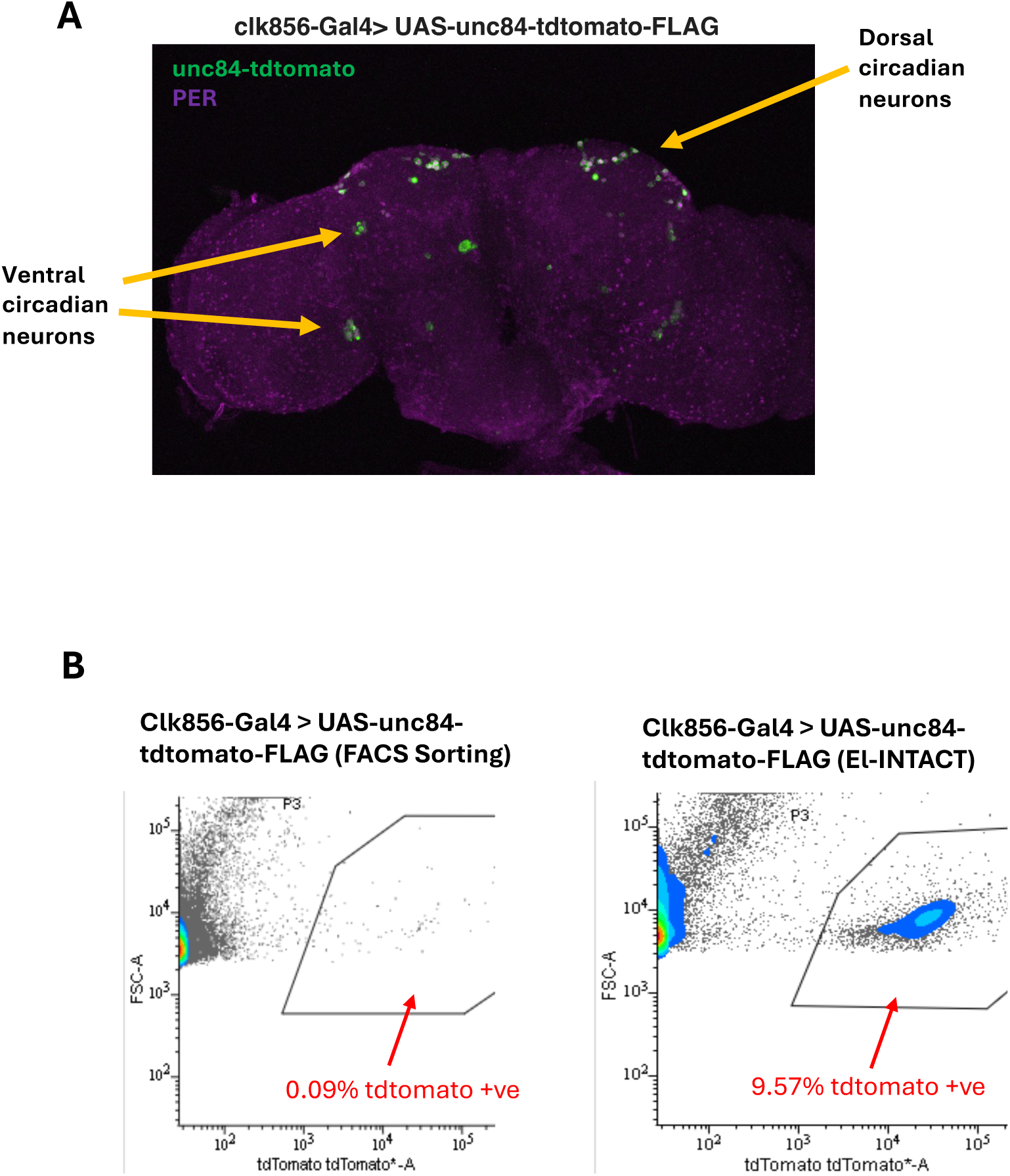
El-INTACT purification of clock neurons using the Clk856-Gal4 driver. A) UAS-unc84-tdtomato-FLAG flies were crossed with Clk856-Gal4 to achieve labeling of the circadian neurons. In green are the circadian neurons visualized by staining for tdtomato. The presence of the core circadian protein PER (in magenta) is used to confirm the identity of clock neurons. B) Representative plots for FACS sorting of El-INTACT purified Clk856-Gal4> UAS-unc84-tdtomato-FLAG nuclei. Gating parameters were set up using non-tdtomato nuclei.

**S3.**
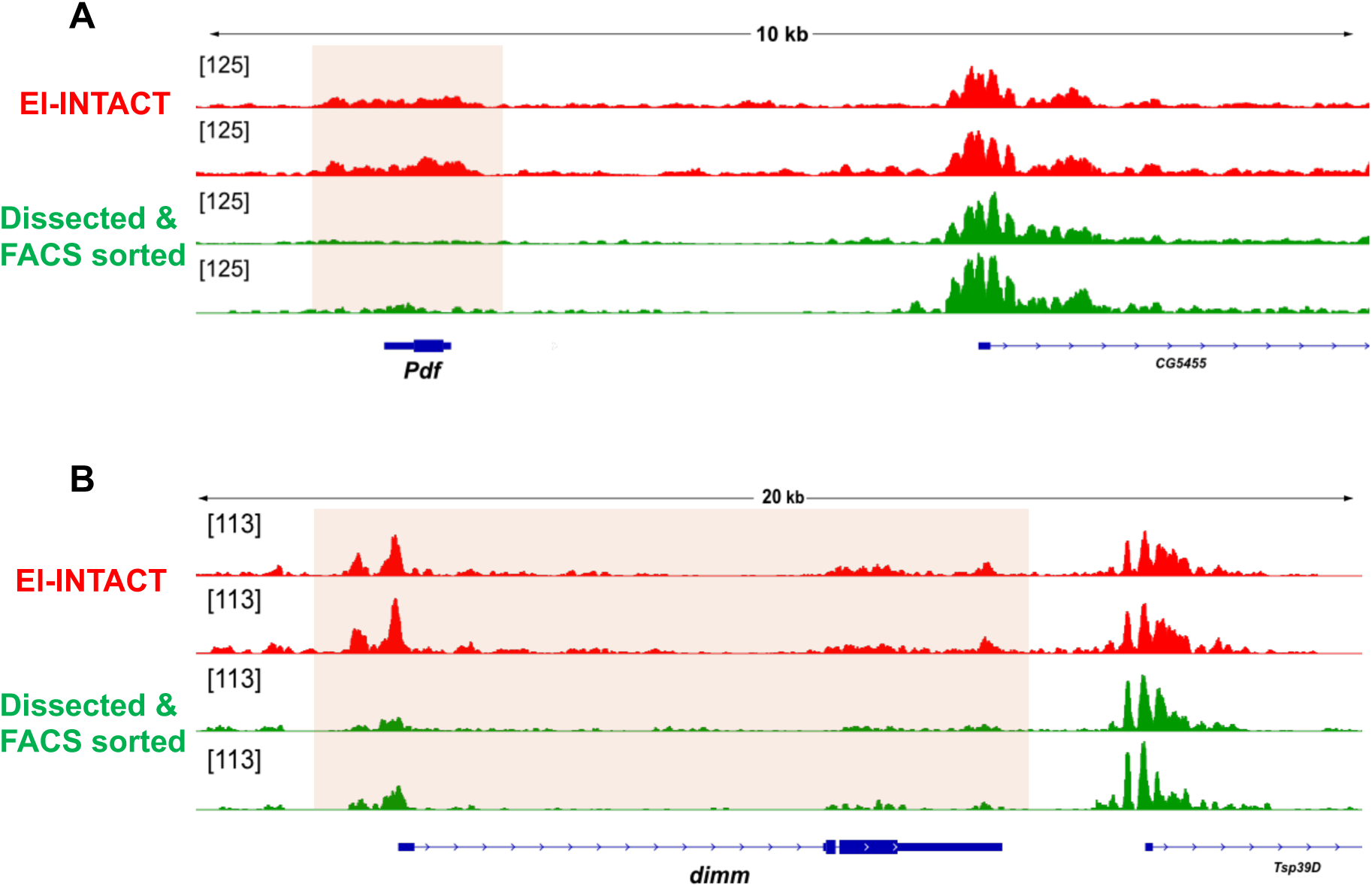
El-INTACT outperforms dissection + FACS sorting to purify difficult to capture cells. ATAC-Seq profiles of clock neurons were generated either using El-INTACT (in red) or dissection and FACS sorting (in green). To make a direct comparison between the different starting material for ATAC-Seq, the same input (4000 cells or nuclei per replicate) was used. Both cells and nuclei were collected at ZT02. Highlighted background regions in the chromatin profiles indicate the genes of interest. A) ATAC-Seq profile of the gene *Pdf* is more accessible in El-INTACT isolated nuclei than in dissected and FACS sorted cells. B) ATAC-Seq profile of the gene *dimm* is more accessible in El-INTACT isolated nuclei than in dissected and FACS sorted cells. Here, chromatin accessibility profile of the neighboring genes CG5455 (A) and Tsp39D (B) serve as internal controls.

**S4.**
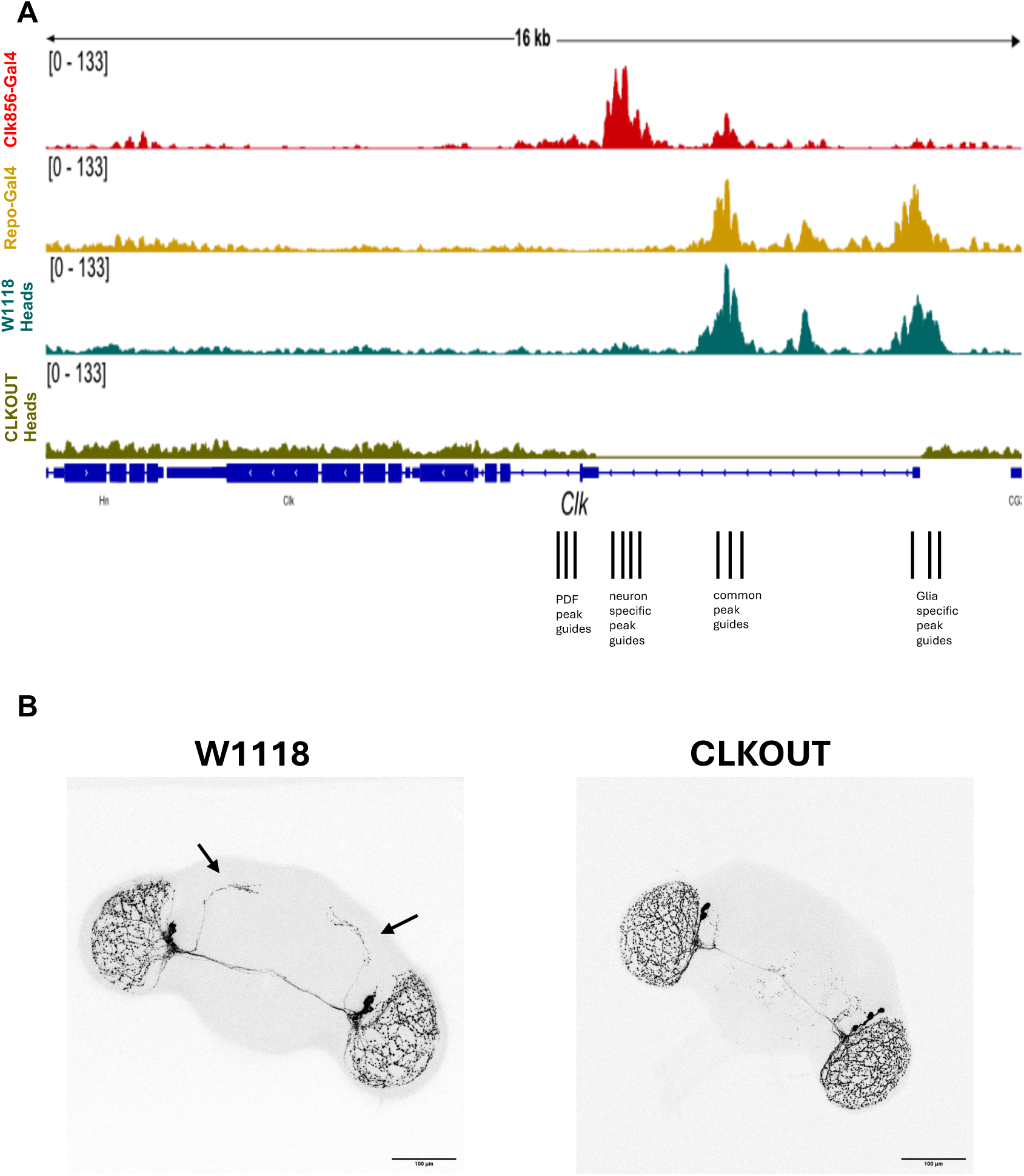
**CLKOUT ATAC-Seq and sLNV projections**A) *Clk* cis-acting region in sorted clock neurons (red), glia (orange), W1118 heads (teal), and ClkOUT heads (olive green). Black lines indicate each guide targeting specific ATAC-Seq peaks. B) Anti-PDF staning (black) in W1118 and CLKOUT flies. Dorsal projections from sLNVs (indicated by black arrows) in W1118 are missing in CLKOUT.

**S5.**
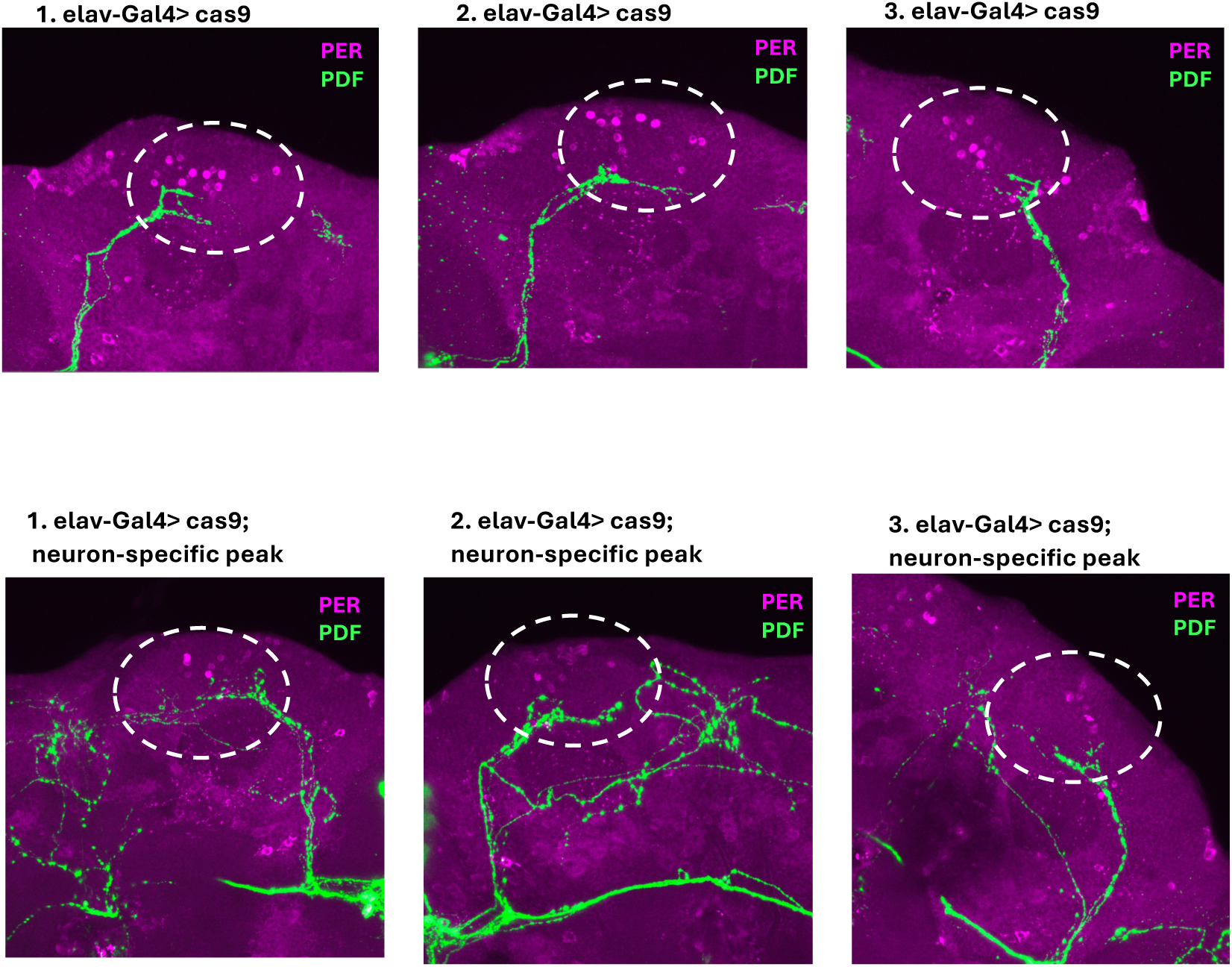
PER expression within DN1ps is also disrupted with neuron-specific guide. PER expression (magenta) in DN1p neurons (indicated by white circle) suggest a significant reduction in PER expressing DN1ps upon neuron-specific peak disruption.

**S6.**
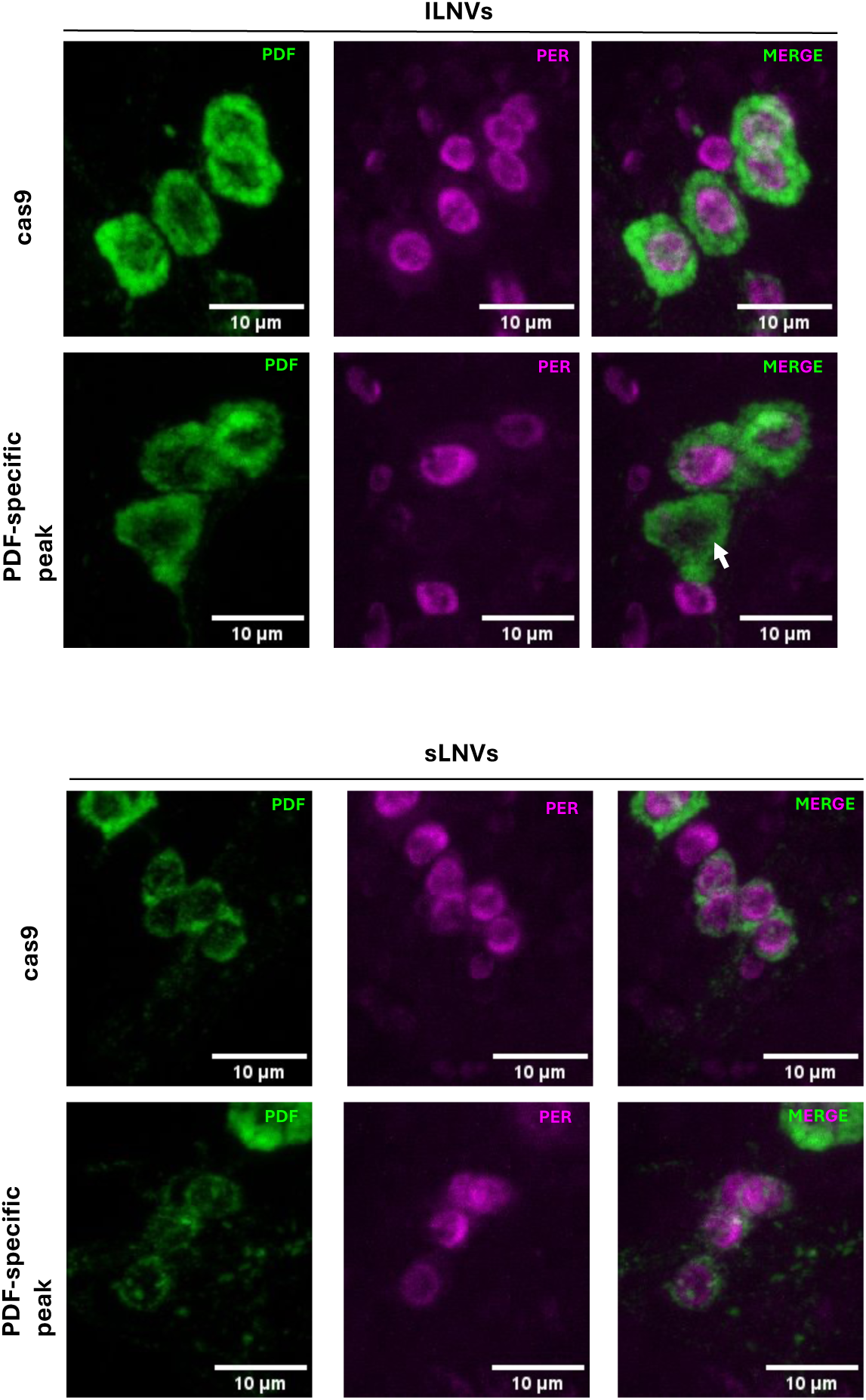
PER expression within the LNVs of PDF peak disrupted flies. PER expression (magenta) in large and small LNVs marked by PDF-expression (green) suggest that PER expression within large LNVs is affected (white arrow) when PDF peak is disrupted. Whereas PER expression in small LNVs remain unaffected.

**S7.**
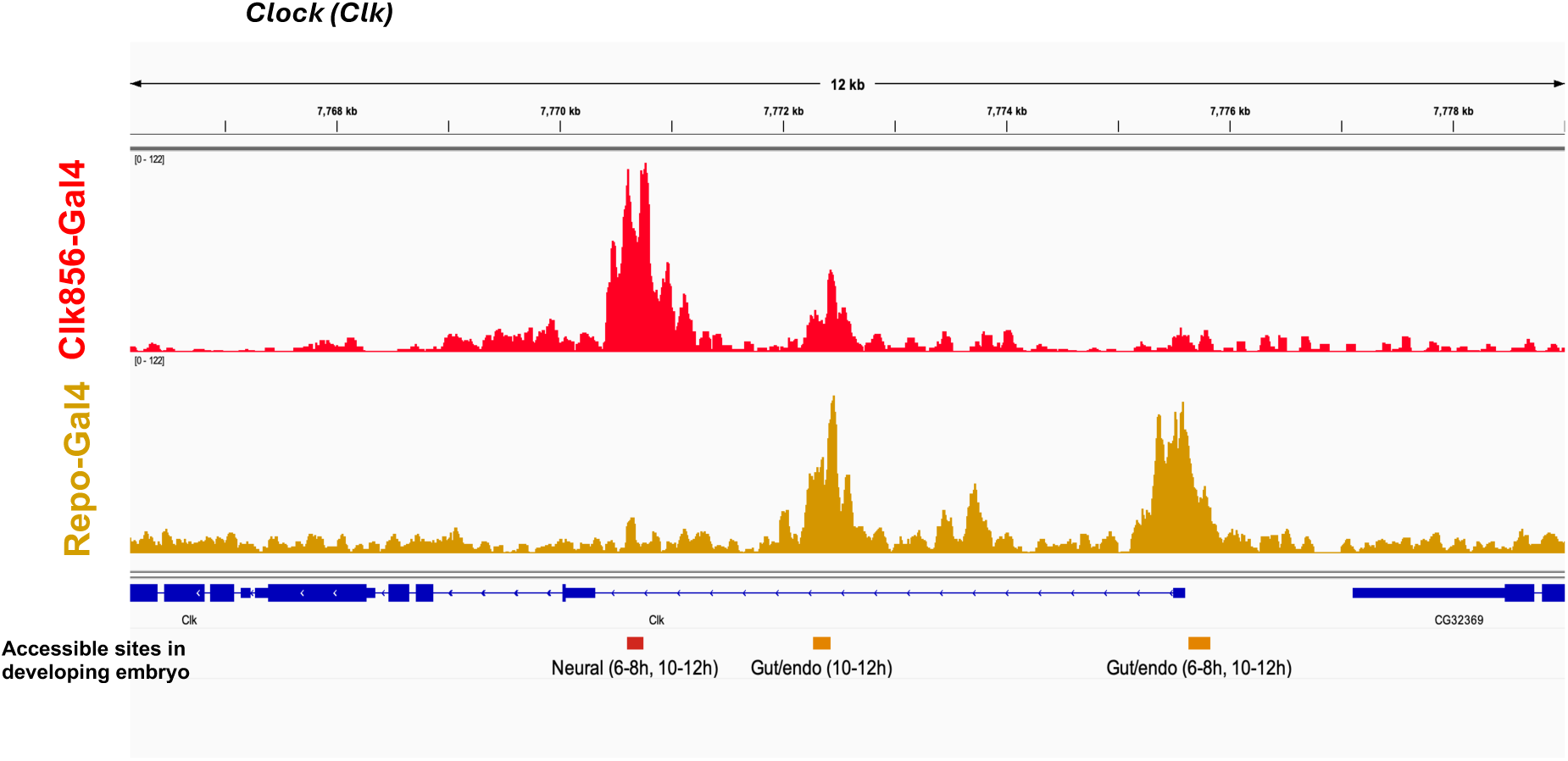
Neuron-specific peak opens up in 6-8 hrs in the developing embryo. Chromatin accessibility across the *D. melanogaster Clk* promoter. Top two tracks: adult bulk ATAC-seq from FACS-purified clock neurons (red) and glia (orange). Third track: embryonic single-cell ATAC-seq peaks that are differentially accessible in a specific germ-layer lineage (“clade”), from Cusanovich et al. 2018 (66), colored and labeled by lineage and by the developmental window(s) in which each peak is clade-specific (2–4 h, 6–8 h, 10–12 h after egg-laying). The neuron-specific adult peak coincides with a Neural-clade (Clade1) embryonic element (red), whereas the two glia-biased peaks coincide with Gut/endoderm-clade (Clade4) elements indicating the clock-neuron core is selectively accessible in the neural lineage from embryogenesis onward.

**S8.**
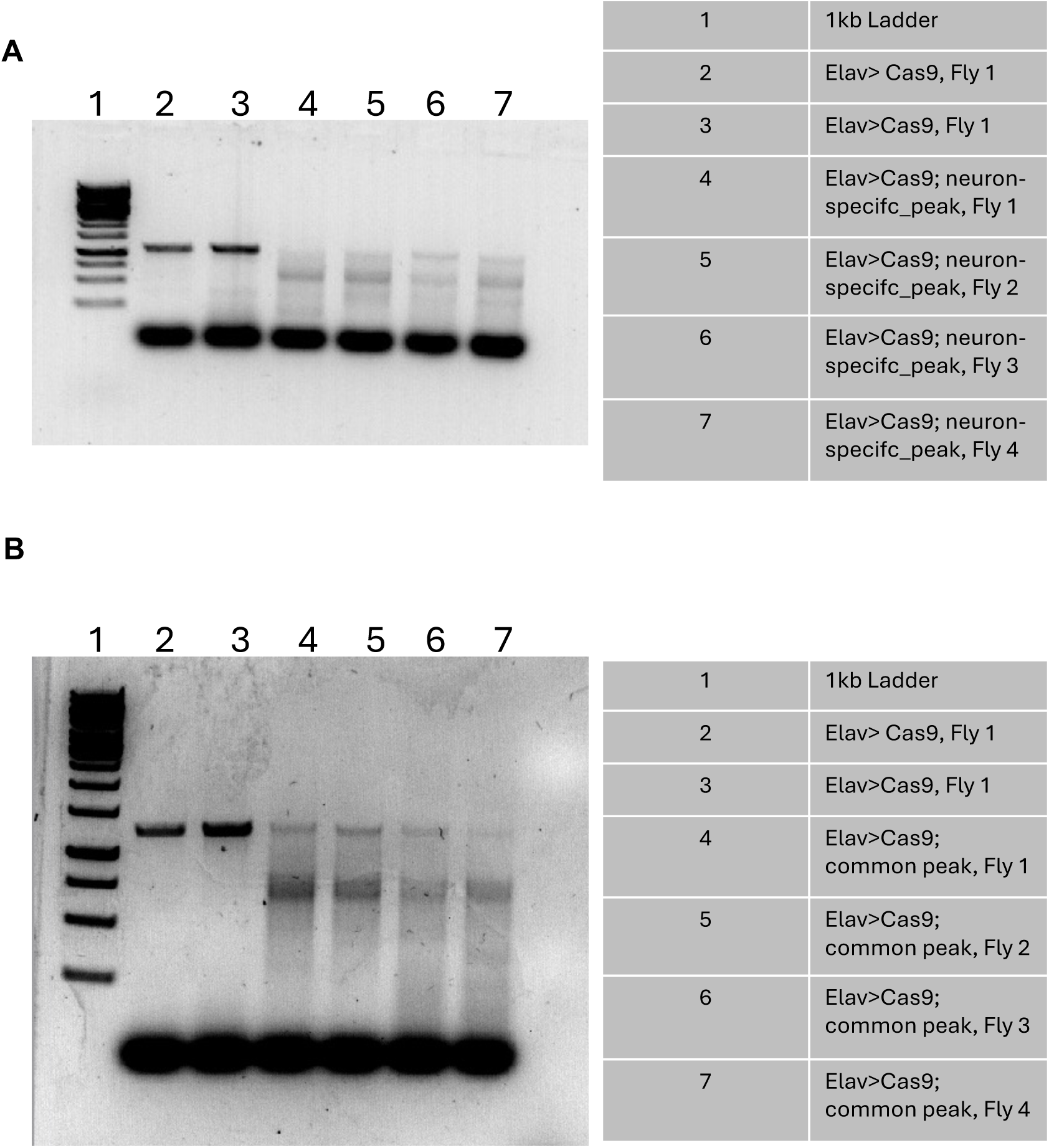
PCR amplification of CRISPR disrupted regions. PCR primers were designed to amplify individual peaks of interest (see Methods). Control lanes yield a single discrete band (lanes 2 and3), whereas mutant lanes 4 -7 show heterogeneous, diffuse products of altered mobility, consistent with a mixed population of indels and deletions at the targeted site. Lower-migrating signal common to all lanes reflects primer dimer.

**S9.**
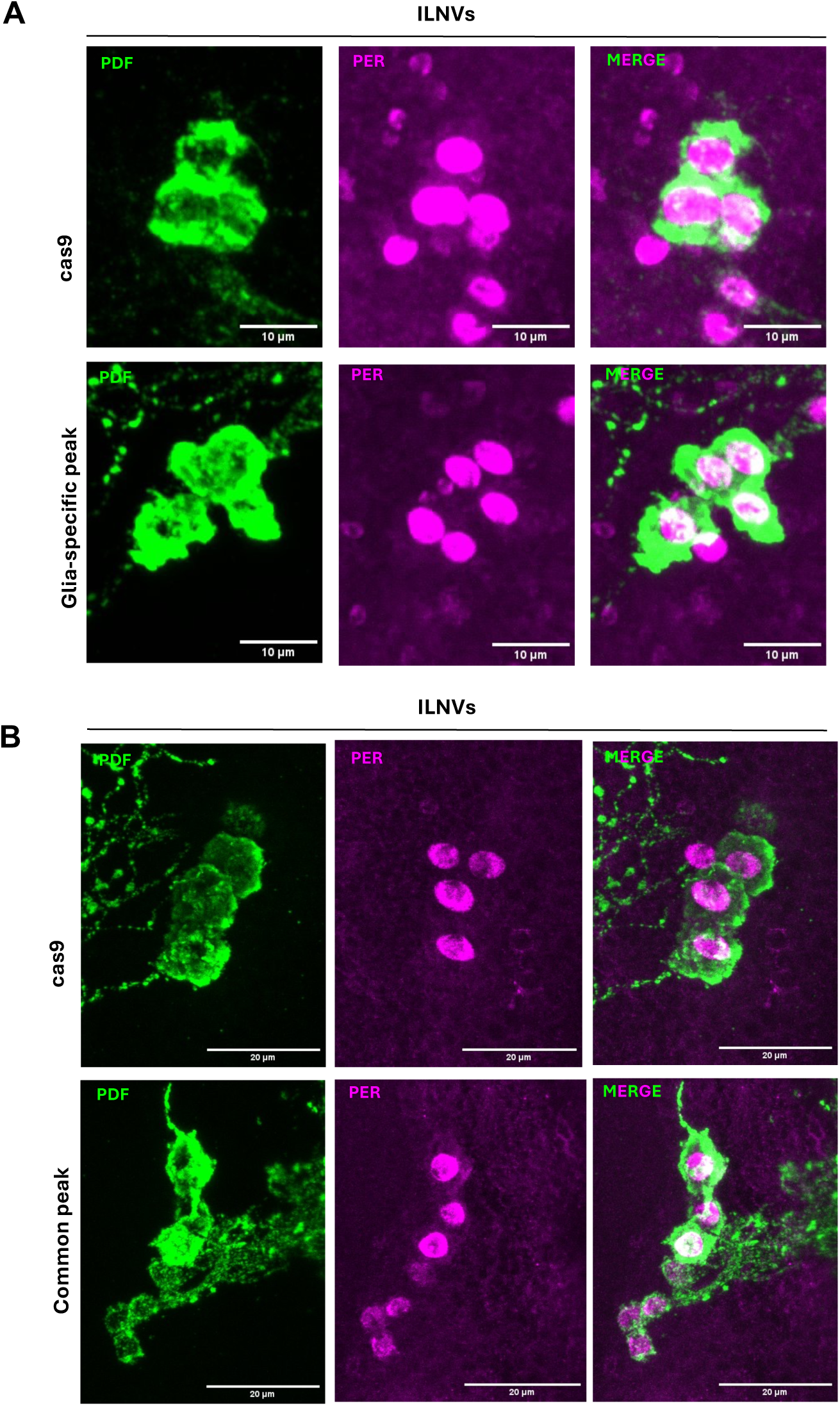
PER expression within the LNVs with common and glia specific peak disruptions. PER expression (magenta) in large a LNVs marked by PDF-expression (green) in glia-specific and common peak disruptions suggest that PER expression within large LNVs remain unaffected.

